# Multi-omic guided iPSC dopaminergic differentiation leads to selective enrichment of PD-vulnerable SOX6+ midbrain neurons with motor recovery of PD models after transplantation

**DOI:** 10.1101/2025.06.18.660098

**Authors:** Roberto Garcia-Swinburn, Guochang Lyu, Judith C. Kreutzmann, Anqi Xiong, Rika Kojima, Carmen Abaurre, Wojciech Paslawski, Katja Schach, Chiara Tremolanti, Per Uhlén, Gonçalo Castelo-Branco, Carmen Salto, Onur Dagliyan, Per Svenningsson, Ernest Arenas

**Author notes:** These authors contributed equally (Roberto Garcia-Swinburn, Guochang Lyu). Deceased 15/09/2024 (Ernest Arenas).

## Abstract

**Background:** Parkinson’s disease (PD) is characterized by the progressive loss of midbrain dopaminergic (mDA) neurons, leading to severe debilitating motor impairment. Recent single-cell analyses have revealed that specific mDA subpopulations, such as SOX6⁺ AGTR1⁺ neurons, are most vulnerable to degeneration. Current human pluripotent stem cell differentiation protocols fail to selectively generate these subtypes.

**Methods:** Here, we describe a multiomic-guided strategy that enriches SOX6⁺ mDA neurons by combining enhancing Sonic Hedgehog (SHH) agonism with prolonged Wnt activation. Initial brute force optimization of a protocol of a prior publication failed to obtain our desired outcome. Using longitudinal single-nuclei (sn) multiomic sequencing, we were able to localize differentiation chokeholds in our timeline and corrected by inducing Wnt more aggressively at the selected time point. After mDA progenitor optimization, we performed *in vitro* validation of downstream mDA differentiation by immunocytochemistry, detection of secreted dopamine by high-performance liquid chromatography and comparison of transcriptomics with publicly available post-mortem midbrain data. Once validated, engineered mDA progenitors were intrastriatally transplanted into hemiparkinsonian mice lesioned with 6-OHDA. Motor recovery was assessed longitudinally, and graft cellular composition was molecularly characterized using immunohistochemistry and snRNA sequencing. Finally, our sequencing dataset was used to validate transcriptional identities of grafted cells compared to postmortem midbrain data, as well as recognize lack of potential detrimental off-targets and pluripotency.

**Results:** Our multiomic approach accelerates floor plate specification, increases expression of mDA developmental markers, and substantially increases the yield of SOX6⁺ mature mDA neurons compared to prior attempts. Following intrastriatal transplantation into hemiparkinsonian mice, these cells restored motor function in approximately 4 months and generated grafts containing mature, stably committed SOX6⁺ A9-like neurons that mirror their endogenous human counterparts.

**Conclusions:** Our work establishes a differentiation platform for generating the PD-susceptible mDA subtype, providing a foundation for precision disease modelling and subtype-targeted cell replacement therapies.

## Background

Parkinson’s disease (PD), the second most prevalent neurodegenerative disorder, lacks curative treatments, severely impacting patients’ quality of life. This condition is characterized by the progressive degeneration of midbrain dopaminergic (mDA) neurons in the substantia nigra pars compacta (SNpc). The loss of these neurons, which project primarily to the dorsal striatum, depletes dopaminergic input and leads to the characteristic motor symptoms of PD [1]. Although advances in pharmacological therapies have been made, these treatments are often insufficient in halting disease progression or restoring lost neuronal function in later stages [2]. Cell therapy offers a promising alternative, with the potential to replace the damaged or lost DA neurons critical to PD pathology [3].

Advances in stem cell and midbrain development research have enabled the generation of transplantable mDA progenitors from human pluripotent stem cells (hPSCs) [4–7]. Success in preclinical models has led to the development of clinical grade mDA progenitors for PD clinical trials [8–11] with recent phase I/II clinical trials reporting safe outcomes, thereby paving the way for future medical use of hPSCs [12–14].

However, the translational promise of hPSC-derived cell therapies hinges on the precise composition and quality of the grafted cells. Single-cell transcriptomic studies have begun to molecularly define the diversity of human mDA neurons [15,16], both in development and in hPSC-derived midbrain cultures [7,16–18]. These studies have revealed a critical challenge: not all mDA neurons are equally susceptible to PD. Recent work has reported complications like dyskinesias, which are linked to serotonergic neuron contamination in fetal transplants [19]. Thus, the inability of pre-existing differentiation protocols, which generate heterogeneous mixes of mDA subtypes, to specifically target this critical population represents a significant limitation for both disease modelling and the development of precision cell therapies.

Single-cell transcriptomics have revealed significant diversity within mDA neurons in the adult human SNpc, categorizing them into three major subtypes based on the expression of *SOX6*, *CALB1*, and *GAD2*, establishing that the SOX6 subpopulation is the most susceptible to degeneration [20,21]. These subtypes emerge during human mDA neuron development [15], and recent single-cell epigenomic profiling (e.g., ATAC-seq) further elucidates midbrain differentiation [22]. To generate PD-vulnerable SOX6^+^ mDA neurons, we first developed a simplified differentiation protocol based on recently identified key developmental regulators [7], but molecular phenotyping revealed unstable phenotypes at later stages, indicating insufficient patterning. This prompted reviewing multiomic data leading to implement two key protocol modifications: enhanced ventralization via stronger Sonic Hedgehog (SHH) pathway induction and prolonged Wnt activation improved midbrain patterning, with elevated *EN1* and *SOX6* expression in engineered cells. Transplanting these cells into hemiparkinsonian (hemiPD) rodent models demonstrated their clinical potential, yielding grafts enriched with SOX6^+^ mDA neurons and restoring motor function. Thus, we here report a multiomic-guided differentiation strategy that specifically enriches for the critical PD-vulnerable mDA subtype and validates its therapeutic potential *in vivo*.

## Materials & Methods

### Cell lines

KOLF 2.1J induced pluripotent stem cell (iPSC) line was obtained through Jackson Laboratories [23]. All experiments performed with these cells followed the Ethical Permit 2021-06727-01 issued by the Swedish Ethical Review Authority. Karyotyping by G-banding, genotypes by polymerase chain reaction (PCR), pluripotency by pluripotency marker microscopy analysis, and mycoplasma infection by PCR were routinely assessed before starting new experiments. iPSC lines were adapted to grow on Laminin 521 (Biolamina, LN521)-coated plates and maintained in Essential 8 Flex media (E8 Flex, Thermo Fisher Scientific) at 37°C, 5% CO_2_, until starting differentiation. The medium was changed daily, and cells were passaged at 70% confluency using TrypLE™ Select Enzyme (Thermo Fisher Scientific). When thawing and replating, 10 μM Rho-associated protein kinase inhibitor Y-27632 (Tocris) was added into the culture media overnight.

### Midbrain dopaminergic neuron differentiation

Two days prior to the mDA differentiation, adapted iPSC cells were seeded at a density of 200,000 cells/cm^2^ onto plates coated with Geltrex (Life Technologies) and Laminin-511 (Biolamina, LN511), and maintained in E8 Flex media (Thermo Fisher Scientific) for expansion. The differentiation protocol was initiated when the cells reached approximately 80% confluency. For the first iteration, cells were cultured at day 0-4 on plates coated with Geltrex and LN511, replated at day 11 on LN511, and replated again at day 16 on LN511 + poly-L-ornithine. Cultured cells were kept in Neurobasal medium (NB) supplemented with N2, B27 (Gibco) and 2 mM L-glutamine (L-Glu; Invitrogen), along with various supplements for the first stage of the protocol. From day 0 to day 4, the culture medium was conditioned with 250 nM LDN193189 (Stemgent), 10 μM SB431542, 1 μM Purmorphamine, and 0.7 μM CHIR99021 from day 0 to day 3, with the inclusion of 10 μM Y27632 (Tocris Bioscience) during the first 24 hours. Boosted CHIR99021 (7.5μM) was administered from day 4 to day 11, while LDN193189, SB431542 and Purmorphamine were withdrawn after day 7 and N2 supplements after day 10. From day 10 to day 28, cells were switched to the differentiation media: Neurobasal/B27/L-Glu media supplemented with 20 ng/mL brain-derived neurotrophic factor (BDNF, R&D Systems), 20 ng/ml glial cell line-derived 455 neurotrophic factor (GDNF; R&D Systems), 1 ng/ml transforming growth factor type b3 (TGFb3; R&D Systems), 0.2 mM ascorbic acid (Sigma), 0.2 mM Dibutyryl cAMP (Sigma). On day 11 and 16, the resulting cells were dissociated into single cells, replated at a density of 800,000 cells/cm2 onto plates coated with LN511 and polyornithine/LN511 respectively, and treated with 10 μM Y27632 for 48 hours. To promote neurogenesis, the mDA differentiation media was supplemented with 10 µM DAPT (Sigma) from day 16. The small molecule GW3965 10 μM was administered from day 12 to day 15, followed by a cocktail of small molecules consisting of 1 μM PD0325901 and 5 μM SU5402 (Sigma), from day 16 to day 21. For the second iteration, conditions were similar except for the following: Culture plates were coated with Geltrex + LN511 until day 14, and poly-L-ornithine + LN511 from day 14 onwards. Cells were only replated once on day 14 and boosted CHIR99021 (7.5μM) was administered from day 4 to day 14. Day 0-7 Purmorphamine concentration was increased to 2 µM, and both PD0325901 and SU5402 were removed from the protocol.

### Immunocytochemistry

Progenitor cells at different stages were seeded in a 96-well plate at a density of about 300,000 cells per well for the intended cell culture purposes. Harvested cells were fixed by 4% paraformaldehyde (PFA) for 20 min at 4°C, and then pre-incubated by 5% donkey serum in phosphate buffered saline containing 0.3% Triton X-100 (PBST) for 1 hour. The samples were incubated with primary antibodies at 4°C overnight. The samples were incubated with either Alexa488 or Alexa555 or Alexa647-conjugated secondary antibodies (Thermo Fisher Scientific) for 30 min and then were incubated with 4’,6-diamidino-2-phenylindole (DAPI, Sigma, D9542) for 15 min. The primary antibodies were used as follows: CORIN (rat, 1:1,000, R&D Systems, MAB2209), DCX (goat, 1:500, SantaCruz, sc-8066), EN1 (mouse 1:50, DSHB, 4G11), FOXA2 (goat, 1:500, R&D Systems, AF2400), GIRK2 (rabbit, 1:400, Alomone, APC006), LMX1A (rabbit, 1:4,000, Millipore, AB10533), LMO3 (goat, 1:200, SantaCruz, sc-82647), MAP2 (mouse 1:1,000, Sigma, M4403), NURR1 (rabbit, 1:500, SantaCruz, sc-990), OTX2 (goat, 1:1,000, R&D Systems, AF1979), TH (rabbit, 1:1,000, Millipore, AB152), TH (mouse, 1:500, ImmunoStar, 22941), and TH (sheep, 1:500, Novus Biological, NB300). Mounted cells were further visualized using confocal microscopy (Zeiss LSM880 or 980) with a 10X or 20X lens. Quantification of images was performed using QuPath [24] by generating a “single measurement classifier” or “composite classifier” for positive or double-positive cells respectively.

### Calcium imaging

The 35mm culture dishes were loaded with the Ca2^+^-sensitive fluorescence indicator Fluo-4/AM (5 µM; Invitrogen, USA) and Pluronic F-127 (0.625%, ThermoFisher Scientific) and incubated for 30 min at 37°C in Krebs-Ringer buffer, containing: NaCl (119 mM), KCl (2.5 mM), NaH2PO4 (1 mM), CaCl2 (2.5 mM), MgCl2 (1.3 mM), HEPES (20 mM) and D-Glucose (11 mM), with pH adjusted to 7.4. Dishes were washed twice and Ca2^+^-measurements were performed in Krebs-Ringer buffer at 37°C on a confocal spinning disk microscope (Nikon CrEST X-Light V3) equipped with a 20x Air lens. Excitation was assessed at 480 nm at a sampling frequency of 1 Hz. The equipment was controlled, and data was acquired using the Nikon NIS-Elements software. FIJI, MATLAB (R2020b, MathWorks, USA) and FluoroSNNAP (https://github.com/tapan-patel/FluoroSNNAP) were used to process and analyze the data.

### High-performance liquid chromatography

mDA progenitors were plated at a density of 200,000 cells/cm2 on Geltrex+Laminin 511-coated 24-well plates on day 14 of the differentiation and cultured until day 28. On the day of media collection, cells were washed with 1 mL PBS and incubated in 200 µL/well of Neurobasal minus phenol red+B27+L-Glutamine for 30 min at 37℃, 5% CO_2_. After incubation, the media was collected and stabilized with 0,1 M perchloric acid. Cells were washed with 1 mL PBS and incubated in 200 µL/well of Neurobasal minus phenol red+B27+L-Glutamine supplemented with 50 mM KCl for 10 min at 37℃, 5% CO_2_. After incubation, the media was collected and stabilized as described [7]. The collected media were centrifuged at 16,000 x g for 10 min at 4°C and then filtered through 0.2 µm nylon membrane inserts by centrifugation at 4,000 x g for 5 min at 4°C. The HPLC-ECD system used was a Dionex Ultimate 3000 series (Dionex, ThermoFisher Scientific, USA) and the injection volume was 20 µL for each sample. Analyte separation was performed on a Dionex C18 reversed-phase MD-150 3.2 mm x 250 mm column (3 µm particle size). Column and analytical cell were kept at 30°C and the first and second analytical cell were set to -100 mV and +300 mV, respectively. The mobile phase was pumped at a flow rate of 0.4 mL/min and consisted of 75 mM monobasic sodium phosphate, 2.2 mM 1-octanesulfonic acid sodium salt, 100 µL/L triethylamine, 25 µM EDTA disodium salt and 10% acetonitrile (v/v), pH 3.0 adjusted with 85% phosphoric acid. Chromatograms were acquired with Chromeleon software (Dionex, ThermoFisher Scientific) over an acquisition time of 55 min. Dopamine concentration was calculated for each sample and divided per minute to calculate dopamine release rate.

### Single cell sequencing libraries preparation and data processing

A total of 5 samples (DIV 11, 16, 28, 42, and 56) from KOLF2.1J mDA neuron cultures generated with the first iteration of the protocol were collected for time-course multiomic analysis (ATAC + RNA sequencing); in addition, 3 samples (DIV 14, 16, and 28) from the second iteration of the protocol were collected for single-nuclei transcriptomic profiling. All 8 samples were prepared with commercially available protocols (10x Genomics) and the resulting Fastqc files were processed through the 10X Genomics Cellranger pipeline [25], while 5 multiomic libraries were aligned to the hg38 reference genome using Cellranger-arc default settings [26]. After quality-control filtering, we obtained high-quality 23,534 cells and then we integrated transcriptomic profiles from both conditions using Harmony [27], enabling a comparative analysis by isolating the mDA neuronal lineage from different datasets. Resulting transcriptomic and epigenomic data from individual samples were combined by projects using the ‘aggr’ function in Cellranger or Cellranger-arc, without normalization applied. Objects produced with Seurat V4.4.0 were employed to accommodate aggregated count matrix [28], with the fragment files specifically integrated into Signac v1.9.0 chromatin assays [29]. Quality metrics for gene expression and peak accessibility, such as mitochondrial ratio, cell cycle score, nucleosome signal, and Transcription Start Site (TSS) enrichment, were thereafter calculated to determine quality control cutoffs. Depending on sample quality, following thresholds were applied for downstream analysis: a maximum of 300,000 and a minimum of 1,000 ATAC counts per cell, a maximum of 25,000 and a minimum of 600 RNA counts, at least 250 genes detected, a maximum nucleosomal signal of 1.5, TSS enrichment of at least 2, and mitochondrial content no higher than 15%. These cutoffs reduced the cell count by less than 20% without significantly affecting clustering downstream. Peak calling was performed with MACS2 v2.2.9 using the aggregated fragments file [30], and those mapped to non-standard chromosomes or overlapping hg38 genome blacklist regions were excluded. DoubletFinder v2.0 was applied to identify doublets in gene expression data by flagging instances with a high resemblance to artificial neighbors in gene expression space [31]. AMULET (ATAC-seq MULtiplet Estimation Tool) was additionally utilized to detect potential multiplets in snATAC-sequencing datasets by enumerating regions with greater than two uniquely aligned reads across the genome [32]. All predicted doublets were removed from the downstream analysis. After merging high-quality cells from the relevant projects, count data was normalized using SCTransform V2 function with regression out of mitochondrial or cell-cycle genes. The top 2,000 most variable genes were initially identified through variance-stabilizing transformation, along with 95% of the most frequent peaks in multiomic datasets, as a part of the significant feature selection process. Subsequent dimensionality reduction was performed using the principal components analysis (PCA), in which the top 30/40 principal components were chosen from the elbow plot to represent most of the diversity of datasets. For snATAC-sequencing datasets, singular value decomposition (SVD) was performed with the top 25 of the Latent Semantic Indexing (LSI) layers to reduce the dimensions of the most common peaks. Notably, the first LSI component was excluded from downstream analysis as it often captures sequencing depth (technical variation) rather than biological variation. Putative clusters were identified by a spiking neural network (SNN) modularity optimization-based clustering algorithm, and the optimal resolution of clustering was determined by iterating over different resolution for the most stable segregation. Finally, the calculated nearest-neighbor graph was projected to two dimensions to visualize the distribution of instances by representing with uniform manifold approximation and projection (UMAP). To accurately identify and interpret cell states *in vitro*, we leveraged combined manual cell type annotation using cluster-specific marker genes [33,34], with automatic annotation methods using reference datasets that characterize first-trimester developing human brain [15,35]. Initially, a label transfer was performed with Celltypist [36] and ‘TransferData’ function in Seurat to automatically classify selected dataset with cell type labels from the reference dataset, based on the similarity of query gene expression profiles to selected anchors in the reference dataset. The identity of each cell will be determined by the highest prediction score, and for stringent comparison, cells with a prediction score below 0.51 will be excluded from downstream analysis. After data trimming and reclustering, a manual cluster-based annotation was performed, using both expression levels of canonical midbrain development marker genes and top 50 upregulated genes identified by the ’FindAllMarkers’ function. These upregulated genes were used for pathway and biological process enrichment analysis via Metascape [34], and the top-ranked terms were used to refine annotations and determine cell subtypes or states. To reconstruct the putative trajectory of mDA differentiation *in vitro*, we employed Monocle3 [37] and scVelo [38,39] to infer pseudo-and latent-temporal ordering of cells using default parameters [40]. The root of trajectory for pseudotime analysis was set as ProgFP or NbM at the earliest timepoint. at the earliest timepoint. To represent quantitatively the cell identity similarities with other published human datasets, we performed Metaneighbor [41]. SCENIC [42] was used to visualize regulatory networks and regulon in our developing cells using AUCell scores.

### Animals

Healthy female adult (8-12 weeks old) C57BL/6N-Rag2Tm1-IL2rgTm1-SirpaNOD/Rj (B6RGS) mice were purchased from Janvier Labs. Groups of 5 animals were housed in ventilated cages under a 12 h light-dark cycle with food (standard chow) and water ad libitum at 20-22°C and 50-55% humidity. Tunnels, cardboard nesting material and chewing wooden bars were used for environmental enrichment. All surgical procedures followed aseptic techniques, and for post-surgical recovery, additional Dietgel Recovery (Scanbur) and food pellets were supplied on the cage floor. Animals were randomly allocated to sham control group (n=6), hemiPD group (n=8) and a grafted group (n=8), ensuring all cages had individuals of each group. Briefly, the exclusion criteria were animals which lost over 10% of their pre-surgical weight after 4 weeks were discarded from the experiment. No animals were discarded based on this criterion. The experimental unit was the individual mouse. All procedures were approved by the local ethical committee (Stockholms djurforsoksetiska namnd) and followed the EU directive 2010/63/EU under the ethical permit of Ernesto Arenas Cases and Sandra Gellhaar (12265-2021; 2-3143/23; 2-3595/24; 2-3147/24).

### 6-OHDA mouse model

B6RGS mice were acclimatized 2-3 weeks before surgery to the animal facility and the researcher and to a recovery diet 2-3 days before surgery. For 6-hydroxidopamine (6-OHDA, Sigma) lesions (or sham), animals were injected with analgesia (Carprofen 5mg/kg + Buprenorphine 0.1 mg/kg) 15 minutes before 4% isoflurane induction. Animals were anesthetized at 1-2% isoflurane in air and injected intracranially (ICR) using a Neurostar Robotic Stereotaxic Frame equipped with a 10 μl syringe with 33G blunt needle (Hamilton) 1 μl of 6-OHDA 3.6 μg/μl in sterile ascorbic acid/saline 0.02% (or sham vehicle) in the right medial forebrain bundle [respect to Bregma: AP -1.2 mm, ML +1.2 mm, DV -5.0 mm] at a flow of 0.2 μl/min, raising the needle 500 μm for 3 minutes. The needle was left inside for 2 additional minutes. Animals were placed in warm cages (28 °C) between 3 days and 1 week for better recovery and rehydrated with Ringer Acetate solution if weight decreased by 5%. 6-OHDA hemiPD was confirmed by amphetamine-induced rotation. Briefly, 4 weeks after the lesion, mice were injected intraperitoneally (IP) with d-amphetamine in saline (5 mg/kg). 10 minutes after injection, the mice were recorded for 30 minutes inside a 15 cm wide cylinder in dark conditions, and the number of net ipsilateral rotations (ipsilateral-contralateral) per minute in active movement was calculated using Ethovision XT 17 (Noldus). Mice are considered hemiPD if the values are >6 rotations/minutes.

### hiPSC transplant

HemiPD-confirmed mice of healthy weight (22-30 g) were randomly selected for ICR hPSC injection in the right striatum [respect to Bregma: AP +0.5 mm, ML +2 mm, DV -3.5 mm]. Analgesia and anesthesia were followed as above. For ICR transplants, 2 µl of 100.000 ± 20.000 hPSCs/µl in Neurobasal medium was injected to achieve 200.000 hPSC transplants. For cell injection, the Neurostar robot was coupled with a 10 µl syringe with a 27G sharp syringe. A flow of 0.5 µl/min was used, and the needle remained for 5 additional minutes. Cells were injected between 1-6 hours after thawing.

### Amphetamines-induced rotations

Mice were analysed a week before surgery, 4 weeks after surgery and hereon once a month until endpoint was reached. Mice were injected IP with 5 mg/kg d-amphetamine in saline (0.5 mg/ml). 10 minutes after injection, the mice were recorded for 30 minutes inside a 15 cm wide cylinder in dark conditions, and the number of net ipsilateral rotations (ipsilateral-contralateral) per minute in active movement (body center traveling at >0.5 cm/s) was calculated using Ethovision XT 17.

### Pole test

Mice were analysed a week before surgery, 4 weeks after surgery and at the endpoint. Animals were placed facing upwards on the top of a 20 cm vertical smooth wooden pole with 2 mm grooves notched every 0.5 cm down. The time to climb down was counted starting when the mouse rotates facing downwards. Animals were trained for a minimum of 2 days before testing day, by performing 3 mock tests each day. If the animals took more than 1 minute to descend, the trial was interrupted and repeated at least 10 minutes later. A total of 3 trials were used to calculate a mean value per animal.

### Cylinder wall

Mice were analysed a week before surgery, 4 weeks after surgery and at the endpoint. Animals were placed inside a 12 cm diameter glass cylinder in red light and quiet conditions. For 15 minutes, animals were recorded and videos were analysed at x8 speed. Unilateral full paw contacts were counted. Paw dragging or paws not fully placed on glass were disregarded.

### Tape removal test

Mice were analysed a week before surgery, 4 weeks after surgery and at the endpoint. A green adhesive paper tape 2x3 mm was placed on each paw individually. The time it took to remove the tape was counted, beginning when the mouse recognized it and ending when it was removed from the paw. Net contralateral time (ipsilateral-contralateral) was calculated to better visualize and eliminate external factors. Before testing day, animals were trained for a minimum of 2 days, performing 3 mock trials for each paw. If the animal took more than a minute, the trial was interrupted and repeated at least 15 minutes later. A total of 3 trials were used to calculate a mean value.

### Immunohistochemistry

5 animals were terminally anesthethized with IP pentobarbital 100 mg/kg dose and then perfused intracardially with 50 ml of pH-corrected PBS1X followed by 50 ml of PFA 4% both at a flow of 9 ml/min. Brains were extracted and placed in PFA overnight, after which they were placed in sucrose 30%/PBS solution at 4°C for 2-3 days. Once cryoprotected, brains were kept at -80°C. Brains were equilibrated at -20°C for 4 hours before slicing. Samples were sliced in series at -20°C at 20 µm using a Cryostar NX70 (Epredia) and stored for further use at -20°C in cryoprotective solution (Ethylenglycol 30%, Glycerol 30%, PBS1X). A total of 5 brains were processed.

Samples were stained using free-floating protocols. Briefly, 2 series of 5 minutes of PBS1X, followed by 15 minutes of PBS-TritonX 0.3% (PBTx0.3%), then 2 series of 5 minutes of PBTx0.1%. Samples were blocked for 1 hour with blocking solution (BSA 1 mg/ml, Normal Donkey Serum 5%, PBTx0.1%) and incubated o/n at 4°C with primary antibody diluted in blocking solution. Primary antibody was washed with 2 series of 5 minutes PBTx0.1% and samples were incubated for 4 hours with secondary antibody diluted in blocking solution and washed twice for 5 minutes with PBS1X. Tissue was counterstained with DAPI 1:1000 in PBS1X for 10 minutes and then washed twice for 5 minutes with PBS1X. Stained samples were mounted on Superfrost Plus slides (Epredia) and Fluoromount-G mounting medium (00-4958-02, Thermo Scientific). Primary antibodies used were STEM121 (mouse, 1:1000, Tanaka Bio, Y40410), TH (rabbit, 1:1000, Pelfreeze, P4010; sheep, 1:250 Pelfreeze, P60101), HuNu (mouse, 1:1000, Millipore, MAB1281), SOX6 (rabbit, 1:200, Abcam, ab30455), ALDH1A (rabbit, 1:500, Atlas Antibodies, HPA002123), AGTR1 (rabbit, 1:100, Invitrogen, MA5-56587), Calbindin-1 (rabbit, 1:200, Invitrogen, PA5-85669), LMX1 (rabbit, 1:100, Millipore, AB10533), FOXA2 (rabbit, 1:200, Cell Signaling, 3143S), NR4A2 (rabbit, 1:500, Invitrogen, PA5-78097), PITX3 (rabbit, 1:100, Invitrogen, 35-2850), DAT (rat, 1:250, Abcam, ab5990), GIRK2 (rabbit, 1:500, Alomone Labs, APC006), VMAT2 (rabbit, 1:500, Sigma-Aldrich, AB1598P), DDC (rabbit, 1:500, Novus Biological, NB300-252), Ki67 (rabbit, 1:200, Abcam, ab15580), SOX2 (goat, 1:200, R&D Systems, AF2018). All fluorescent secondary antibodies were from Invitrogen, donkey as host and conjugated with Alexa Fluor 488, 555 or 647. The HRP-conjugated antibody was a goat anti-mouse from Invitrogen. Images were captured either with laser confocal microscope Zeiss LSM800 equipped with x20 lens or high-throughput epifluorescence microscope Zeiss Axioscan equipped with x10 lens. Quantification of images was performed using QuPath [24] by generating a ‘single measurement classifier’ or ‘composite classifier’ for positive or double-positive cells respectively.

### Single-nuclei RNA sequencing of grafts and data analysis

3 animals were euthanized by cervical dislocation, and brains were promptly extracted and flash-frozen in dry-ice chilled isopentane for 10-15 seconds. Flash-frozen brains were stored at -80 °C until graft extraction. A total of 3 brains with grafts were processed independently.

On the day of nuclei extraction, brains were equilibrated to -20°C and sliced in -20°C cooled chambers. Brain matrices were used to stabilize the brain, and 1 mm-thick slices for a total of 4 (2 before the center of graft implant location and 2 after) were cut using chilled razors. 1 mm punches of each slice of the dorsal striatum were extracted using 1 mm biopsy-punchers (49101, PFM Medical).

Nuclei were dissociated using the 10X Genomics Chromium nuclei extraction kit (1000494) using manufacturer’s instructions. For better extraction, eppendorfs used for tissue dissociation were coated with BSA 1% o/n at 4°C, briefly washed with filtered sterile PBS1X before procedure. 10X Genomics Chromium GEM-X Single Cell 3’ Kit v4 (1000691), Chromium GEM-X Single Cell 3’ Chip Kit v4 (1000690) and Dual Index Kit TT Set A (1000215) were used for GEM generation using manufacturer’s instructions with the 10X Controller iX. GEMs were sent for RNA-sequencing to National Genomics Infrastructure of SciLifeLab (Karolinska Institute, Sweden) using the NovaSeq XPlus 25B. Data analysis was performed as the *in vitro* RNAseq dataset, described above. To ensure analysis of human cells, we aligned the libraries to the h38 human genome dataset and the m39 mouse genome dataset, and cells with a ratio of human:mouse gene count >1 were processed.

### Statistics

The N for the animals was calculated using G-Power 3.1 [43] based on prior data from several publications of hemiPD rodent model behavior [44]. Briefly, using the “ANOVA: repeated measures, between factors” a priori statistical test, the effect size was calculated to be 0.82, α =0.05, β=0.95, groups =3, measurements=7 and correlation among repeated measurements=0.8. GraphPad Prism v10.00 was used for generating all graphs and performing statistical analyses. For pairwise comparisons, Two-tailed Student’s t-test was utilized. One-way ANOVA or two-way ANOVA with Tukey’s or Bonferroni’s multiple comparison tests were used to compare multiple variables. Error bars indicate the standard error of the mean (SEM). F are reported with degrees of freedom for numerator and denominator (F_DFn,DFd_). Significance levels were set as follows: ns = not significant; * = p<0.05; ** = p<0.01; *** = p<0.001; **** = p<0.0001.

## Results

### Longitudinal multi-omic single-nuclei sequencing delineates cell composition throughout the process of midbrain dopaminergic neuron differentiation *in vitro*

Building upon our recently published method [45], we designed a new protocol focused on precise midbrain floor plate (mFP) patterning and improving cell viability to generate sufficient progenitor cells. Our protocol produced sequential differentiation of human induced pluripotent stem cells (hiPSCs) into midbrain progenitors, positive in mFP progenitor markers such as FOXA2, OTX2 and LMX1A, and functional neurons expressing canonical mDA markers like TH or Nurr1 (Fig. 1a, Extended Fig. 1).

**Fig. 1:**
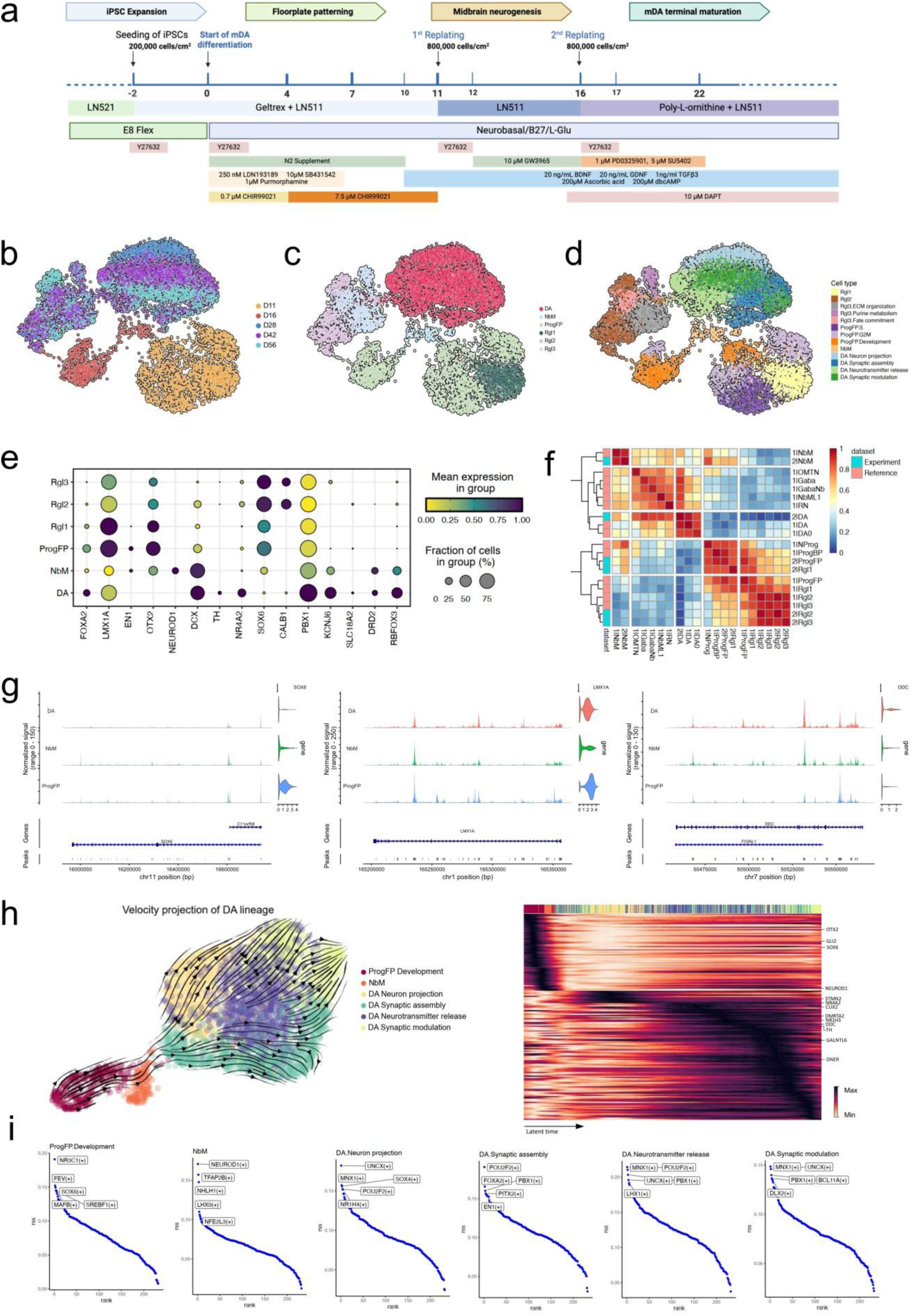
Single nuclei multi-omic sequencing revealed the diversity of cellular composition during mDA neuron differentiation *in vitro.* **a,** Schematic diagram of the initial differentiation protocol used on the iPSC. **b,** UMAP of time-course samples profiled with simultaneous snRNA-seq and snATAC-seq showing clustering of cells based on both transcriptomic and chromatin accessibility profiles. **c,** Annotated bimodal UMAP clusterings based on major cell types. **d,** Subsets of cell types were manually annotated with top-ranked marker genes and their enriched GO terms. **e,** Dot plot visualizing the expression levels and the percentage of cells expressing curated key genes across cell types during mDA neuron differentiation. **f,** Heatmap of Area Under the Receiver Operating Characteristic Curve (AUROC) scores comparing annotated cell types derived from in vitro differentiation in this study with those from a publicly available first-trimester human midbrain dataset. **g,** Normalized chromatin accessibility tracks at genomic loci of genes of interest (SOX6, LMX1A and DDC), along with enriched accession peaks and links. **h,** UMAP clustering showing RNA velocity projection in mDA-related cell types (left), and a heatmap displaying gene expression dynamics of cluster driver genes ordered by their inferred latent time (right). **i,** Regulon specificity score (RSS) plots highlighting the top five cell type-specific transcription factors involved in mDA differentiation.

To delineate developmental trajectories, we performed single-nuclei multiomic sequencing (snATAC-seq + snRNAseq) in benchmarked wild-type iPSCs (KOLF2.1J) at five developmental stages (Days *in vitro* (DIV): DIV11, 16, 28, 42 and 56; Fig. 1b). Following quality control and filtering, we annotated 9,233 high-quality nuclei using integrated manual-automatic classification based on cluster-specific markers and developing human brain references [15] categorized into major types, with differentiation to a mature DA identity based on their enriched Gene Ontology (GO) terms (Fig. 1c,d). Chronologically, early stages (DIV11-16) showed high expression of regional specification markers (LMX1A, OTX2). By DIV28, dopaminergic commitment marker NR4A2 upregulated, while late stages (DIV42-56) exhibited elevated expression of maturation markers (such as TH, NR4A2, KCNJ6), indicating functional specification (Fig. 1e, Extended Fig. 2). Further chromatin accessibility analysis showed precursor cell types such as floor plate progenitors (ProgFP), medial neuroblasts (NbM) and DA clusters to recapitulate expected chromatin access (*SOX6* and *LMX1A* for mDA progenitors, and *DDC* for mature functional mDA neurons) with ProgFP having a higher potential to differentiate to SOX6 populations (Fig. 1g). We then validated our annotations by comparing them to publicly available single cell RNA-seq datasets from human fetal midbrain samples [15], by quantifying cell type identity across individual cells. As a result, our annotated cell types closely resembled their endogenous counterparts, as demonstrated by hierarchical clustering (Fig. 1f). High similarity scores (>0.8) were predominantly observed during early stages of the midbrain differentiation and were primarily enriched in ProgFP and NbM, underscoring the effectiveness of our approach in recapitulating endogenous early mDA neuron development. Radial glia cell subtypes, Rgl2 and Rgl3, also displayed transcriptomic profiles akin to their counterparts *in vivo*, hinting at the establishment of an mDA signaling niche to deliver cell type-specific developmental factors for neuronal maturation *in vitro* [46]. At advanced stages, mDA neurons closely matched their endogenous cells from reference datasets (reference DA and DA0 vs experimental DA similarity score >0.8). These findings show that while mDA neurons matched reference profiles, later stages may benefit from extended maturation cues to enhance postmitotic neuronal survival.

### Reconstructed differentiation trajectory resolves transcriptional dynamics during mDA lineage specification and maturation

To define the molecular logic of *in vitro* differentiation, we reconstructed the developmental trajectory of the dopaminergic lineage. We isolated progenitor and neuronal populations (ProgFP, NbM and mDA neurons) from our multiomic dataset and inferred their relationships using pseudotime ordering and RNA velocity. Both analyses confirmed a direct progression from ProgFP and NbM progenitors to mature mDA neurons, validating the protocol’s ability to recapitulate this fundamental lineage path (Fig. 1h).

Uniform manifold approximation and projection (UMAP) analysis revealed a continuous maturation trajectory for mDA neurons. GO analysis of genes ordered along this path showed a developmentally relevant progression of biological processes: initial neurite outgrowth was followed by neurotransmitter release, synaptic assembly, and finally synaptic modulation (Extended Fig. 2b), matching the pattern observed in pseudotime analysis. While most cells followed the canonical path, a subset of late-stage cells (DIV42–56) appeared to bypass the axonal development phase characteristic of DIV28, suggesting the existence of alternative maturation mechanisms, potentially influenced by identity cues established earlier in development [47]. We next applied single-cell regulatory network inference and clustering (SCENIC) to identify stage-specific regulons. Clustering of AUCell scores for activities of each regulon revealed distinct segregation between early neural progenitors, ProgFP, and NbM, and their descendant mDA neurons along the development trajectory (Fig. 1i). We identified key transcription factors (TFs) orchestrating each developmental transition. In ProgFP cells, the NR3C1 (glucocorticoid receptor) regulon was upregulated, promoting progenitor pool expansion by skewing cells toward proliferation over differentiation [48]. In parallel, SREBF1, a pro-neural transcription factor controlling mDA neurogenesis [49], drove the subsequent transition to neurogenic progenitors (NProg). In NbM cells, pro-neural TFs NEUROD1 and NHLH1 emerged as top regulons, initiating neurogenesis and differentiation in postmitotic cells [50,51]. Subtypes of mDA neurons shared a core set of transcription factors, including PBX1, a canonical regulator of mDA specification and survival [52]. Furthermore, transcription factors such as POU2F2 and BCL11A displayed more specialized roles at distinct maturation stages [18,53]. In conclusion, trajectory reconstruction confirms that our differentiation protocol recapitulates key embryological processes of mDA development. More importantly, this approach provides a powerful *de novo* framework for identifying the developmental regulatory modules, offering deeper insights into dopaminergic neuron differentiation and maturation.

### Enhancing ventralization improves floogrplate patterning and accelerates neuronal conversion

Given that robust regional patterning underpins high-quality mDA neuron generation, and our single-cell data suggested suboptimal ventralization, we hypothesized that enhanced mFP ventralization would improve patterning outcomes. We induced Engrailed 1 (EN1), a well-established marker for mDA neuron specification [54], while streamlining the protocol for scalability. EN1 plays a pivotal role from the earliest stages of regional specification and maintenance of mDA neurons in adulthood [55]. It supports the midbrain-hindbrain boundary formation during development and sustains mDA function by regulating key genes (*TH, VMAT2, DAT,* and *ALDH1A1*) in postmitotic neurons [55,56]. Thus, EN1 serves as both a checkpoint for enhanced mDA neuron specification quality and a predictor of transplantation outcome [9].

To enhance EN1 expression, we applied a higher concentration of the SHH agonist purmorphamine [6], and optimized culture conditions to maintain cell viability (Fig. 2a). Subsequently, elevated purmorphamine significantly boosted EN1 expression at DIV11 (Optimized: 88.53% ± 6.8%; Original: 20.00% ± 3.65%) without altering core floor plate markers FOXA2 or LMX1A (Fig. 2b,e; Extended Fig. 3a). The effects of this enhanced ventralization were more pronounced at later stages. By DIV16, during the fate transition from NbM towards early mDA neurons, new protocol cultures exhibited an accelerated TH^+^ neuron generation (Optimized: 26.12% ± 3.93%; Original: 3.57% ± 1.68%) and increased co-expression of TH^+^, EN1^+^ and SOX6^+^ (Optimized: 75.92% ± 1.26%; Original: 7.56% ± 5.11%; Fig. 2c,f). This improved molecular signature persisted through maturation (DIV28), with sustained co-expression of EN1 and SOX6 (Optimized: 65.44% ± 7.06%; Original: 5.35% ± 3.75%; Fig. 2d,g, Extended Fig. 3b). These neurons exhibited functional maturity, displaying spontaneous DA release (0,19 pmol/min), and increased DA release rate by KCl induction (0,48 pmol/min) by DIV28 (Fig, 2h). Additionally, calcium dynamics characteristic of mDA neurons such as spontaneous activity and responsiveness to KCl was also observed by DIV28 (Fig. 2i). All these data combined suggest that the enhanced ventralization of mDA progenitors produces functional mDA neurons as early as DIV28. To molecularly characterize the improvements driven by enhanced ventralization, we performed single-nuclei RNA sequencing (snRNA-seq) on cells from the new protocol at DIV14, 16 and 28. The new protocol accelerated differentiation, with a higher proportion of NProg and NbM emerging at earlier stages, accompanied by a reduction in ProgFP cells (Fig. 3b). Using the same annotation methods as in the previous dataset, the resulting mDA lineage cell types were enriched with classic mDA neuron marker genes (Fig. 3c), including *FOXA2, LMX1A, OTX2, TH,* and *NR4A2*, and showed an expected developmental trajectory across different cell populations (Fig. 3d). Critically, postmitotic cells exhibited higher EN1, sustained OTX2 and SOX6, and elevated maturation markers TH, SLC18A2 (VMAT2), and CALB1 [57] (Fig. 3f). SCENIC analysis identified the key transcription factors activated by the new protocol (Fig. 3e). The paired-box transcription factor PAX7 became the dominant regulon in NProg, suggesting precise rostro-caudal patterning as it contributes to establish the midbrain-hindbrain boundary during regionalization [58,59]. Other leading regulons in NProg and NbM cells are also actively involved in guiding the neuronal identity acquisition. For instance, POU3F1 and ZEB1 are essential regulators of neural lineage commitment, often working synergistically to coordinate correct orders of neuronal development, as POU3F1 promotes early neural fate decisions, while ZEB1 modulates the developmental timing and prevents premature differentiation [54,60]. Notably, PITX3 emerged as a leading regulon in mDA neurons, which reflects the progressive acquisition of mature mDA phenotypes [61] and highlights their molecular similarity to endogenous SNpc mDA neurons (Fig. 3e,f). Regulon activity profiles from the integrated dataset revealed a distinct and significant regulatory impacts of TCF4 and TCF12 under the new protocol (Extended Fig. 4c). Interestingly, the new protocol also appeared to enhance SREBF1 expression throughout differentiation, linking it to its known role in regulating midbrain dopaminergic neurogenesis [49] , and its involvement in controlling neuronal dendrite expansion through lipid homeostasis [62]. Overall, a targeted enhanced ventralization, guided by multiomic analysis, improved the specification of mDA neurons and drove them toward a transcriptomic profile that closely resembles that of vulnerable SNpc neurons.

**Fig. 2:**
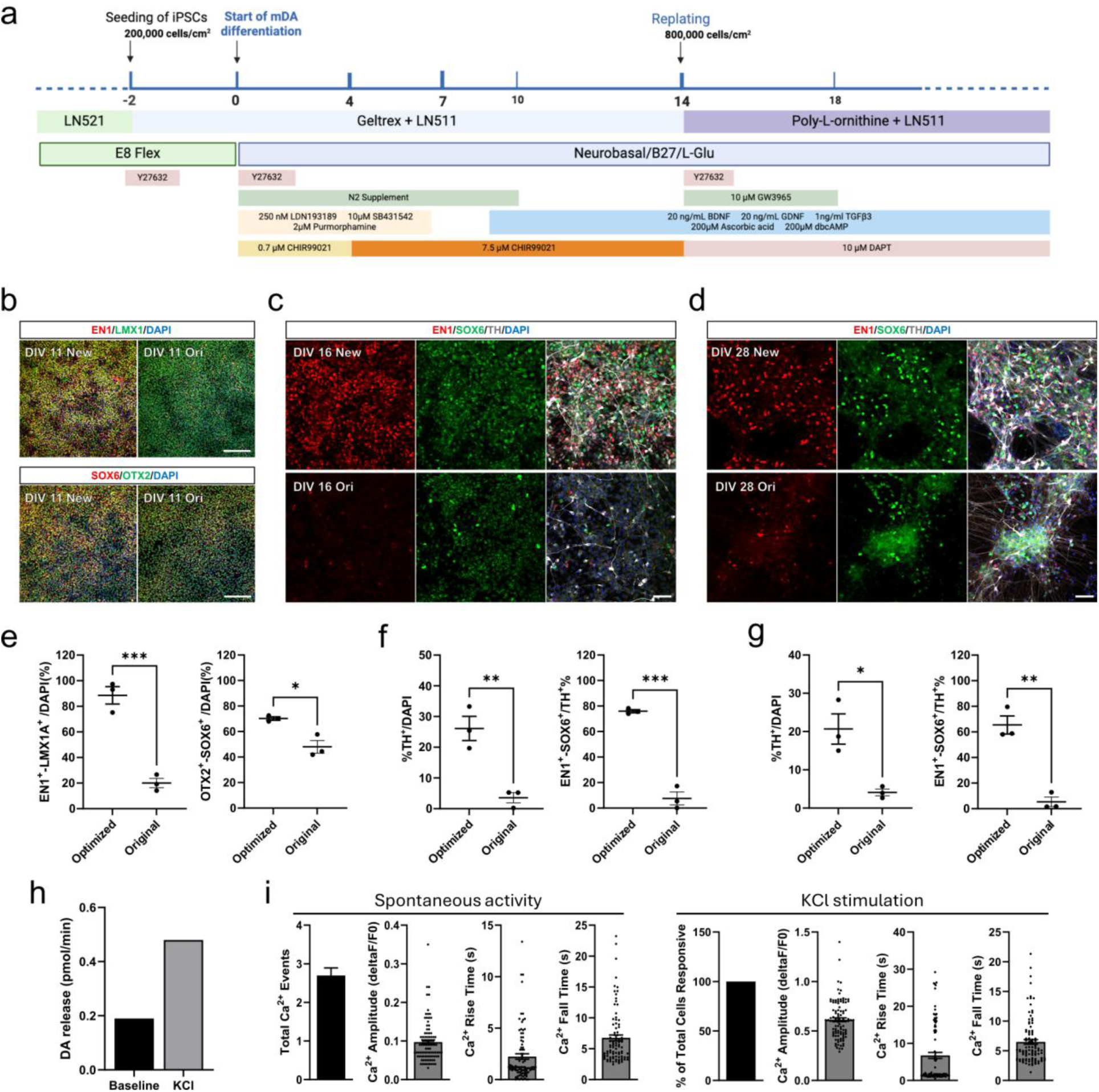
Prolonged ventralization promotes accelerated differentiation of mDA neurons. **a**, Schematic of the optimized differentiation protocol, illustrating the application of pharmacological compounds and growth factors. Representative immunocytochemistry images and quantification from both original protocol and new protocol showing **b,e**, efficacies of midbrain floorplate patterning at DIV11, showing co-localizations of LMX1^+^/EN1^+^ and SOX6^+^/OTX2^+^; and EN1^+^/SOX6^+^/TH^+^ neurons at **c,f,** DIV 16 and **d,g,** 28. White scale bars = 1 mm in B, 50 µm in **c**,**d** (n = 3 biological replicates). **h**, Dopamine release analyzed by HPLC of media of DIV28 of optimized protocol after 30 minutes of spontaneous release (baseline) and 10 minutes of KCl-induction (KCl). **i**, Calcium imaging recordings of spontaneous and KCl-induced activities on DIV 28 of optimized protocol, presenting the total number of calcium events per neuron recorded, and the percentages of cells responding to KCl-induced depolarization. Metrics of activities, including amplitude, rise time, and fall time of calcium signals, are depicted in the subsequent bar plots. Each dot represents one cell. Two-tailed T-tests in **e-g** were performed. * = p<0.05; ** = p<0.01; *** = p<0.001. Error bars = SEM.

**Fig. 3:**
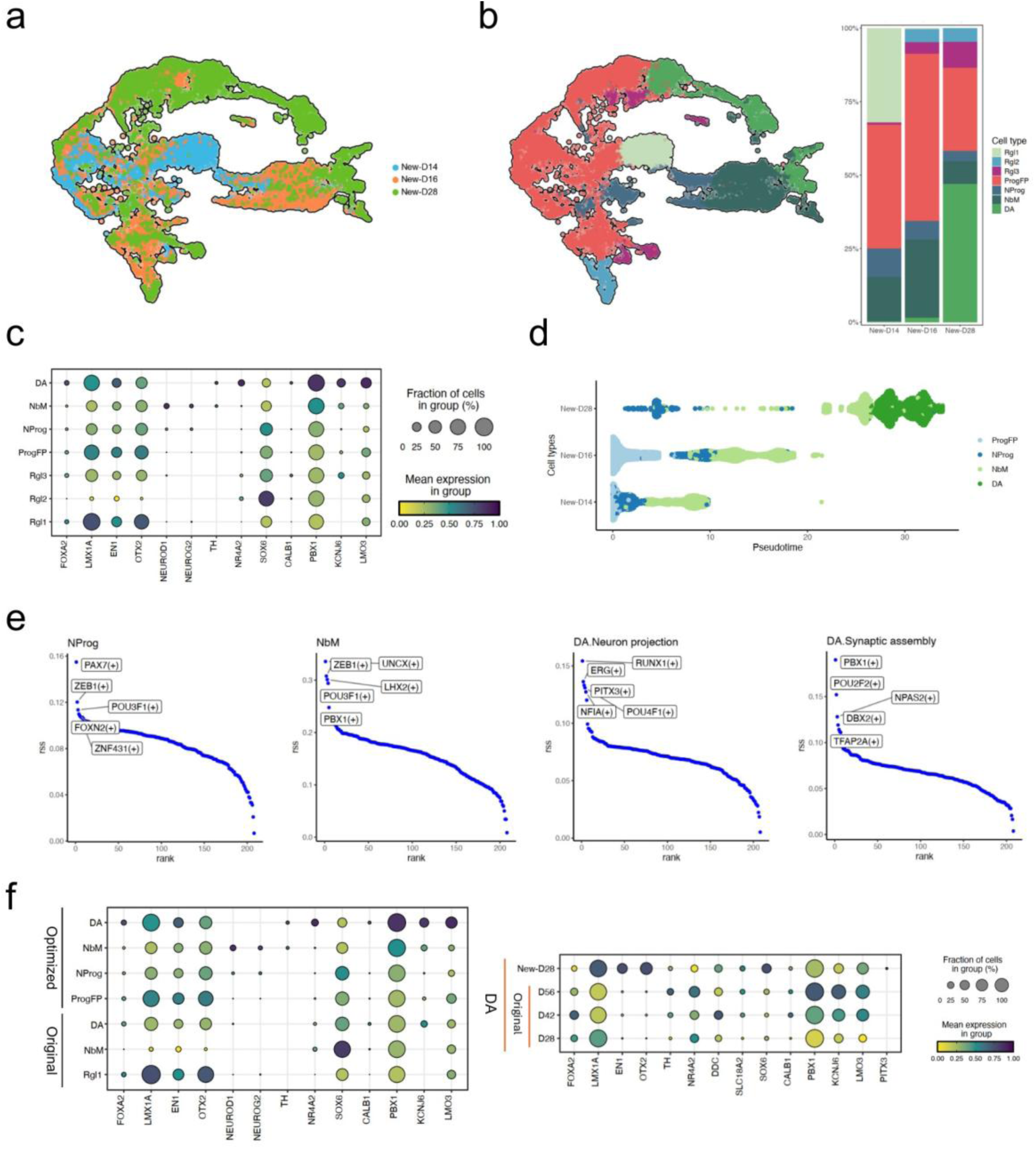
snRNA-seq confirms enhanced molecular patterning under the optimized protocol. a, UMAP embeddings of mDA differentiation in vitro using KOLF2.1J iPSCs under the new protocol, color coded by different time points at DIV 14, 16 and 28. **b,** UMAP embeddings of detailed cell types were manually annotated using top-ranked marker genes and their enriched GO terms, and proportion of cells classified into each cell type per differentiation time point. **c,** Dot plot visualizing the expression levels and the percentage of cells expressing curated key genes across distinct cell types generated under the new protocol. **d**, Pseudotime trajectory analysis showing progression from progenitor-like populations (ProgFP, NProg, and NbM) to differentiated DA neurons. Cells are ordered by inferred pseudotime along the x-axis. **e**, Regulon specificity score (RSS) plots highlighting the top 5 cell type-specific transcription factors driving mDA differentiation under the new protocol. (f) Dot plots showing the expression levels and the percentage of cells expressing midbrain dopaminergic lineage marker genes after integration in (left) midbrain cells categorized as dopaminergic neurons (DA) (right), dopaminergic lineage cell types across all timepoints and both conditions.

### Transplanted iPSC-derived mDA progenitors mature into striatum-innervating dopaminergic neurons and restore motor function in hemiPD mice

Having established that our optimized protocol enriches ventralized mDA subpopulation *in vitro*, we next evaluated the therapeutic potential of our engineered cells *in vivo* using hemiparkinsonian rodent models. We unilaterally lesioned immunocompromised B6RGS mice by injecting 6-OHDA into the medial forebrain bundle. Upon confirming hemiparkinsonism, we intrastriatally transplanted 200,000 cells into randomly selected mice. Motor recovery was assessed longitudinally by amphetamine-induced ipsilateral rotations, showing that grafted hemiPD animals returned to pre-lesion state 4 months post-implant (m.p.i.), which was maintained 1 month later, ending our grafting experiment (Sham: -0.24 ± 0.73; 6-OHDA: 13.41 ± 1.43; Grafted: -0.70 ±1.18 net ipsilateral rotations/min). To ensure overall motor recovery, we also analysed complex motor coordination by tape removal test (Sham: -0.40 ± 0.34; 6-OHDA: 19.51 ± 2.50; Grafted: -2.82 ±0.96 net ipsilateral removal time) and pole test (Sham: 5.00 ± 0.59; 6-OHDA: 21.25 ± 3.53; Grafted: 8.56 ± 1.13 seconds), and spontaneous motor activity by cylinder wall test (Sham: 51.82 ± 3.73%; 6-OHDA: 8.85 ± 4.75%; Grafted: 40.89 ± 3.73% ipsilateral contacts). Human cells were detected in these 5-month grafts by STEM121^+^ staining, confirming that grafted cells did not migrate to other regions (Fig. 4c). Furthermore, human DANs were detected by double positive staining of TH and HuNu, with these cells showing extended processes, characteristic of mDA neurons (Fig. 4d,e). Using stereological quantification of TH^+^/HuNu^+^ cells (Fig. 4F), grafts were estimated to have 160000 human nuclei (155700 ± 35893), of which about 10% were also stained positively for TH (10.36 ± 0.73%). Additionally, no actively proliferative human cells were found within the grafts (Extended Fig. 5)

**Fig. 4:**
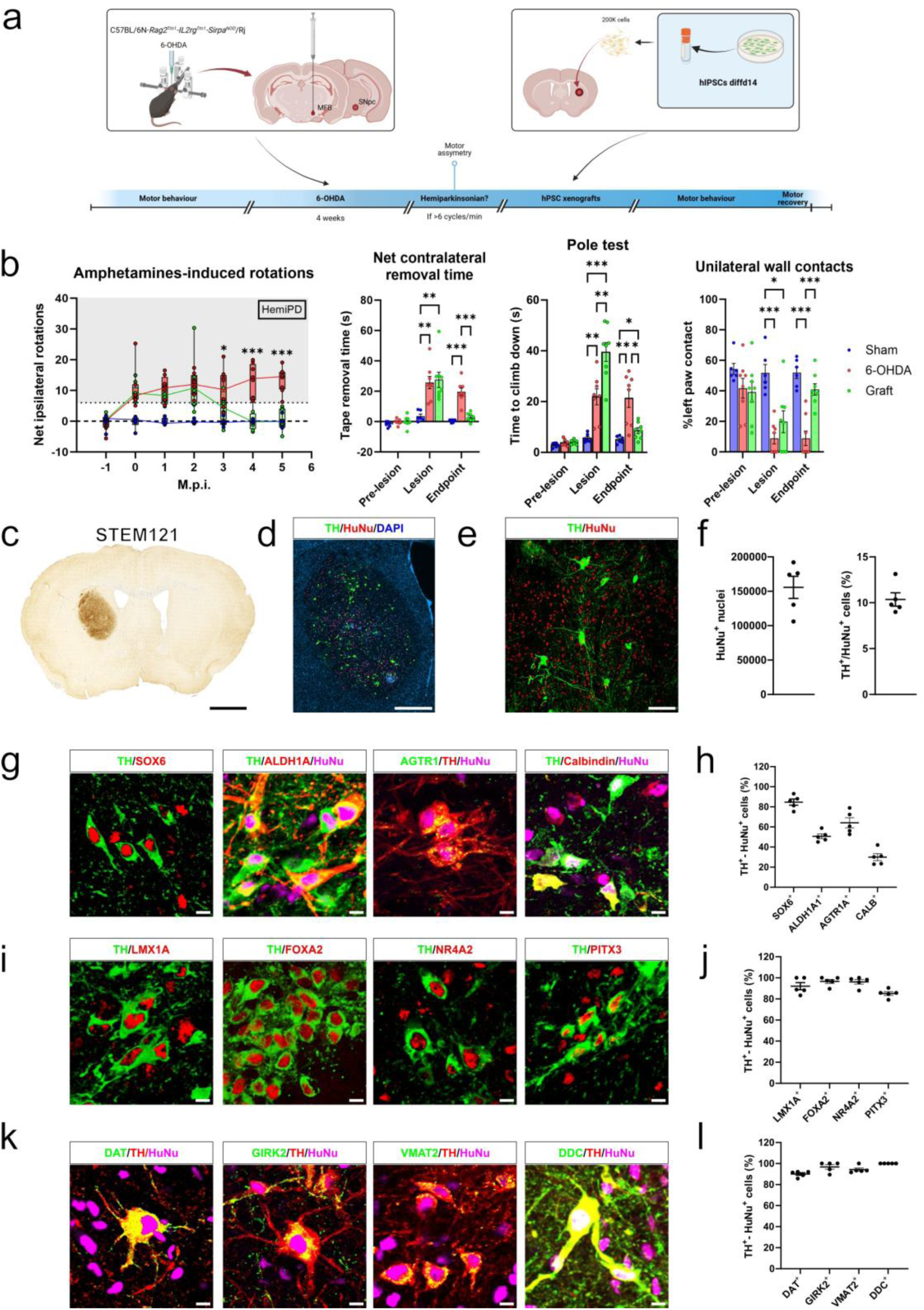
Transplanted hiPSC-derived mDA progenitors restore motor function and mature into A9-subtype mDA neurons *in vivo*. **a**, Diagram of the experimental design for the implantation of day 14-differentiated hiPSC in B6RGS mice. **b**, Motor recovery is observed by continuous analysis of amphetamine-induced ipsilateral rotation, endpoint tape removal test, endpoint pole test and endpoint cylinder wall analysis (right). Of note, grafted animals recover pre-lesion state while the 6-OHDA control animals maintain the lesioned state. **c**, Example of grafted tissue, stained against for STEM121, showing the human cells within the graft. **d,** Micrograph of graft stained against for TH and HuNu, showing hDANs all over the graft. **e,** Closeup of grafted hDANs showing elongated and ramified extensions. **f,** Stereological estimate of HuNu+ nuclei (left) and percentage of hDANs (TH+-HuNu+) in grafts (right). **g,** Representative examples of hDANs stained positive for A9-subtype marker, SOX6, ALDH1A1, AGTR1A and A10 marker, Calbindin. **h,** Quantification of percentage of hDANs (TH+-HuNu+) expressing SOX6, ALDH1A1, AGTR1A and Calbindin. **i,** Representative examples of hDANs stained positive for mDA neuron makers, LMX1A, FOXA2, NR4A2 and PITX3. **j,** Quantification of percentage of hDANs (TH+-HuNu+) expressing LMX1A, FOXA2, NR4A2 and PITX3. **k,** Representative examples of hDANs stained positive for proteins involved in DA excretion, DAT, GIRK2, VMAT2 and DDC. **l,** Quantification of percentage of hDANs (TH+-HuNu+) expressing DAT, GIRK2, VMAT2 and DDC. m.p.i = months post-implant. n for animal behavioral experiment **b** = sham: 6; 6-OHDA: 8; hIPSC-grafted: 8. N for IHC **f,h,j,l**, = 5. Scale bar = **c**: 1mm; d: 500 µm; **e**: 100 µm; **g,i,k**: 10 µm. Means are represented as bars (**b**) or lines (**f,h,j,l**). One-way ANOVA with Tukey correction between treatments was performed separately on each timepoint in the continuous amphetamine test (F_12,114_ = 7.876), as well as for each separate timepoint in the endpoint experiments (Pole test F_4,38,_ = 27.32; Tape Removal F_4.38_ = 8.40; Cylinder Wall F_4,38_ = 3.97). * = p<0.05; ** = p<0.01; *** = p<0.001. Error bars = SEM.

After establishing the presence of dopaminergic neurons (DANs) within our grafts, we aimed to analyse if these DANs recapitulated the phenotypical aspects of the most PD-vulnerable portion of nigral DANs which have been reported to be A9-subtype DANs from the SNpc, more specifically the SOX6^+^/AGTR1a^+^ subpopulation located in the ventral SNpc [20,63]. To this aim, we performed a series of IHC analysis of the grafted DANs. A majority of grafted TH^+^ cells were found to be positive for several markers distinctive to the PD-vulnerable A9-subtype (Fig. 4g,h), such as SOX6 (84.67 ± 3.03%), ALDH1A1 (50.67 ± 2.37%) and AGTR1 (64.16 ± 4.97%). In contrast, a minority of the DANS stained positively for the A10-subtype marker Calbindin (29.92 ± 3.45%). Another aspect that we wanted to evaluate is the expression of canonical markers for mDA neurons in the grafted TH^+^ cells (Fig. 4i,j). We performed IHC and we identified approximately all of the grafted DANs as LMX1A^+^ (92.09 ± 3.37%), FOXA2^+^ (96.66 ± 1.88%), NR4A2^+^ (96.19 ± 2.16%) and PITX3^+^ (85.14 ± 1.80%). Finally, we performed another series of IHC to evaluate expression of proteins involved in DA regulation and secretion (Fig. 4k,l). We were able to determine that the grafted DANs expressed DAT (90.03 ± 1.23%), GIRK2 (96.78 ± 2.12%), VMAT (94.24 ± 1.47%) and DDC (100.00 ± 0.00%). Overall, these results show that grafted cells mature into a large percentage of DANs expressing canonical markers for A9-subtype mDA neurons that are able to ameliorate motor degeneration in rodent PD models.

### snRNA-seq reveals A9-subtype mDA neuron enrichment in grafts

To unbiasedly characterize graft composition and transcriptional fidelity to the transplanted neurons to native human neuron subtypes, we performed snRNA-seq on graft tissue from three transplanted mice. After quality control, we retained 9196 high-quality human nuclei. Unsupervised clustering revealed that the graft composition consisted of four main cell lineages: neurons, radial glia, gliobasts and oligodendroglial lineage cells, with no presence of vascular leptomeningeal meningeal cells (VLMC) contamination, an off-target population in mDA neuron differentiation [64] (Fig. 5a, Extended Fig. 6). Further analysis of the neuronal clusters (defined by RBFOX3) resolved into four distinct populations: dopaminergic (DAergic), glutamatergic, GABAergic, and a small cluster of neural progenitors (defined by high expression of INA and low expression of distinctive neuronal markers) (Extended Fig. 7a, Extended Fig. 8a).

**Fig. 5:**
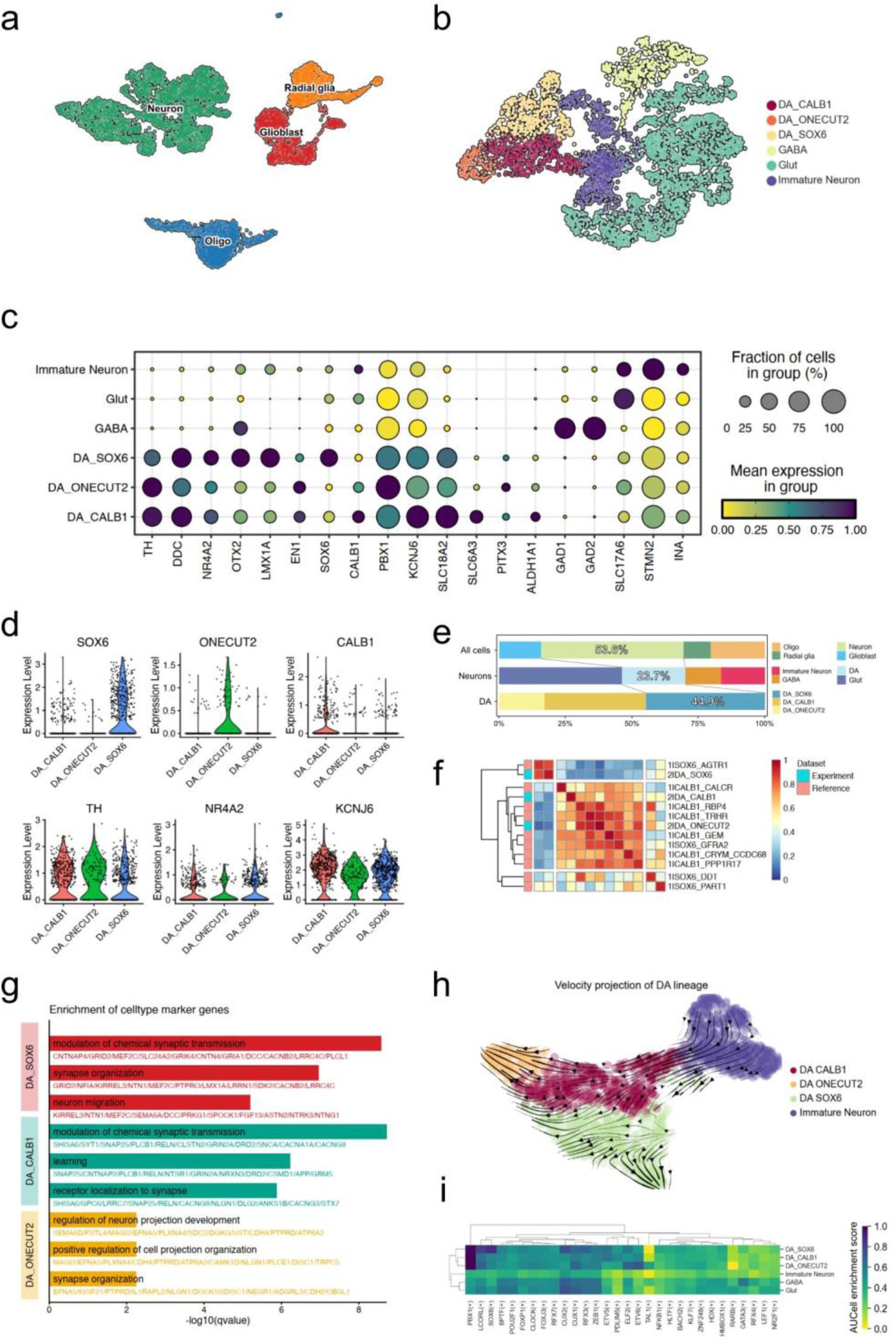
snRNA-seq of grafts reveals presence of transcriptionally-defined A9-subtype neurons. **a**, Annotated UMAP embeddings of 9,196 high-quality human nuclei recovered from three grafted mice at five months post-transplantation, with cells classified into neurons, radial glia, gliobasts and oligodendrocyte precursors; and **b**, neuronal population was consisted of immature neurons, GABAergic neurons, Glutamatergic neurons and 3 subtypes of mDA neurons, each characterized by specific transcription factors, as DA_CALB1, DA_ONECUT2, and DA_SOX6. **c,** Dot plot showing average gene expression and percentage of expressing cells for selected marker genes across annotated neuronal cell types. **d**, Violin plots of representative marker genes for dopaminergic subtypes (SOX6, ONECUT2, CALB1), pan-DA markers (TH, NR4A2, KCNJ6), illustrating subtype-specific and shared expression patterns. **e**, Average proportion of yielded human nuclei in each cellular hierarchy across three individual datasets, demonstrating consistent representation of DA subtypes (N=3). **f**, MetaNeighbor analysis between transplanted mDA neurons in this study and a reference dataset of DA neurons from the SNpc region of PD patients and matched controls, suggesting transcriptional similarities between DA_SOX6 subtype and SOX6_AGTR1 subtype from the reference dataset. **g,** GO enrichment analysis of DA subtype-specific markers, showing distinct biological processes enriched in DA_SOX6, DA_CALB1, and DA_ONECUT2 populations. **h**, RNA velocity projection of mDA neuron trajectories in UMAP embeddings, suggesting differentiation pathways among different DA subtypes and immature neurons. **i**, Heatmap of average regulon activities of the top 30 transcription factors across mDA and other neuronal subtypes.

Critically, the DAergic cluster was comprised of 3 distinct subclusters: DA_SOX6, DA_CALB1 and DA_ONECUT2 (Fig. 5b). DA clusters exhibited high expression of canonical dopaminergic markers (*TH, DDC, NR4A2, LMX1A, PBX1, SLC18A2* [VMAT2], *SLC6A3* [DAT], *PITX3, KCNJ6* [GIRK2]), whereas GABA and glutamatergic clusters lacked these but expressed *GAD1* and *GAD2* and *SLC17A6* [VGLUT2], respectively. INA clusters were distinguished from glutamatergic clusters by elevated *SLC17A6*, *STMN2* and *INA* expression (Extended Fig. 7b,c, Extended Fig. 8b,c). Within the DA lineage, subclusters were resolved by differential expression of SOX6, CALB1 and ONECUT2 (Fig. 5c,d). Roughly half of DA cells were classified as DA_SOX6 (44.73 ± 4.79%), with the remainder predominantly DA_CALB1 and a minority DA_ONECUT2 (Fig. 5e). These results confirm our protocol’s capacity to produce PD-susceptible SOX6⁺ mDA subtype *in vivo*.

### Transcriptomic comparison validates fidelity of grafted neurons to native human midbrain subtypes

To rigorously benchmark the identity of our grafted dopaminergic neurons, we compared their transcriptional profiles directly to those of native adult human mDA cells. We performed a MetaNeighbor analysis, a quantitative method for assessing cell type similarity, comparing our graft-derived DA subclusters to the well-annotated dopaminergic subtypes from a reference single cell atlas of the human substantia nigra [20] (Fig. 5f). This analysis showed high distinctive resemblance with a score >0.8 between our DA_SOX6 subcluster and the SOX6_AGTR1 subcluster from Kamath dataset, which have been reported to be the most susceptible subpopulation to PD [20,63]. In contrast, the DA_ONECUT2 subcluster showed resemblance to the human CALB1_TRHR reported to be the most resilient. However, the DA_CALB1 subcluster lacked exclusive alignment with any Kamath cluster, instead showing broad similarity to multiple CALB1^+^ subtypes, indicating incomplete subtype commitment or ongoing differentiation in this population. These results confirm successful *in vivo* differentiation into SOX6^+^ mDA neurons mirroring PD-vulnerable human counterparts reinforces that we were successfully able to generate cells that differentiate *in vivo* to cells resembling human SOX6 mDA neurons.

### Gene regulatory network analysis confirms stable commitment of SOX6^+^ dopaminergic neurons

Given the transcriptional similarity of grafted DA neurons to native human subtypes, we assessed whether SOX6^+^ subtype represented a stably committed fate or a transitional state. GO analysis revealed distinct functional specializations among the dopaminergic subclusters, consistent with advanced maturation (Fig. 5g). The DA_SOX6 population was enriched for terms related to “synaptic transmission” and “organization”, hallmarks of functional, integrated neurons. In contrast, the DA_CALB1 and DA_ONECUT2 clusters were either associated as not as enriched or more associated with terms like “learning”, suggesting incomplete ventral direction or functional roles more akin to dorsal SN or ventral tegmental area. (Fig. 5g).

To directly infer developmental potential, we performed RNA velocity analysis on our DA and immature neurons subclusters. We found that immature neurons have the potential to become DA_CALB1 cells. Interestingly, the DA_CALB1 subclusters have the potential to become either DA_ONECUT2 or DA_SOX6, which in contrast their velocity show that these identities were a stable endpoint, not a transient phase (Fig. 5h, Extended Fig. 8d,e).

Further evaluation of the gene regulatory network by SCENIC established that the DA clusters were enriched in exclusive TFs expression profile, not found in the immature neuron, GABA or Glut clusters (Fig. 5i, Extended Fig. 8f). For example, PBX1 regulon networks have been associated with ventral midbrain DA organization and are highly enriched (score >0.8) in our DA cluster. Moreover, the DA_SOX6 is differentially highly enriched in SOX6 (score >0.8) and CUX2 (score >0.6) within the DA clusters. Interestingly, the CUX2 regulon network has been reported to be highly associated with the SOX6^+^AGTR1^+^ clusters of neurons in human datasets, further validated in PD macaque models [20,65]. In conclusion, this multi-faceted computational analysis demonstrates that our grafts contain mature, synaptically active neurons whose subtype identities are locked in by stable gene regulatory networks. The DA_SOX6 population represents a terminal, committed fate, confirming that our protocol generates the precise, stable, and vulnerable neuronal subtype required for accurate disease modeling and effective cell therapy for PD.

## Discussion

This study establishes a multiomic-guided framework for differentiating human iPSCs into mDA neurons with high molecular and functional fidelity to the PD-vulnerable SOX6⁺AGTR1⁺ subtype of the substantia nigra. By leveraging single-nuclei transcriptomic and epigenomic data to identify and overcome insufficient ventral patterning, we developed an optimized protocol that achieves accelerated production of these clinically relevant neurons. Furthermore, we were able to reproduce this protocol with different researchers and in different laboratories, addressing reproducibility caveats because of different experiences and/or facilities.

Our longitudinal multiomic atlas of *in vitro* mDA differentiation revealed stage-specific chromatin accessibility and transcriptional shifts mirroring human midbrain development. Late maturation (DIV42-56) coincided with elevated TH, DDC and KCNJ6 expression, and PBX1 regulon activity, a known regulator of mDA survival and functional maturation. Crucially, ventralization enhancement (via strategic agonism of SHH signaling) boosted EN1 expression, a key regulator of midbrain identity and a known predictor of transplant success, yielding ventral SNpc neurons with enhanced molecular fidelity to human native counterparts. Furthermore, we were able to identify regulatory networks involving mDA neuron maturation enriched in the new protocol, such as TCF12 and TCF4. Specifically, TCF12 plays a dual role in the early commitment of neural progenitors and the late specification of neuronal subsets, with its loss leading to altered expression of TH and AHD2 (ALDH1A1), the latter being crucial for the specification and maintenance of dopaminergic neurons, particularly in the A9 dopaminergic neuronal group [66]. TCF4, on the other hand, is likely involved in the fine-tuning of neuron identity during the onset of mDA neuron subtype specification. In a mouse study, deletion of TCF4 led to delayed maturation of dopaminergic neurons, primarily through transient suppression of *Ahd2* and *Nxph3* [67]. A key advancement in the optimized protocol was the precise modulation of midbrain floor plate (mFP) patterning by the biphasic WNT activation (CHIR boost) and SSH agonism, combined with optimized coating conditions using LN511, which generated a substantial population of mFP progenitors by day 11 and enhanced the expression of critical mDA markers such as FOXA2, LMX1A, and OTX2. This improved patterning was accompanied by a reduced contamination by non-neural cells (a common issue in region-specific neuron generation). Furthermore, the specification to committed progenitors allowed us to omit the MEK/ERK inhibitor PD0325901 and the FGF receptor inhibitor SU5402, previously used to suppress expansion of SOX2^+^ progenitors [68], therefore harnessing the intrinsic neuronal differentiation mechanisms without any further external influence for producing mature mDA neurons downstream. The addition of synthetic LXR ligands (GW3965) further accelerated neurogenesis, leading to the earlier appearance of TH⁺ neurons at day 16 compared to day 21 in prior protocols. This earlier establishment enables functional assays (such as calcium imaging and neurotransmitter release analysis) to be performed sooner, thereby expediting detection of phenotypic differences. As an example, live-cell calcium recordings confirmed that the derived mDA neurons displayed spontaneous electrical activity and DA release and robust depolarization responses, consistent with functionally active neurons [69]. Finally, as a proof-of concept validation for therapeutic approaches, we determined that the new protocol could produce mFP progenitors able to differentiate *in vivo* to mDA neurons resembling the A9-subtype nigral neurons in majority, based on expression of proteins such as SOX6, AGTR1, GIRK2 or ALDH1A, reported to be more susceptible to PD in human and animal models [20,63,70,71]. Furthermore, evaluation of the transcriptomic profile of the DA neurons in the graft showed high resemblance to the human counterpart, specially the SOX6-AGTR1A mDA neurons. Moreover, the lack of proliferative progenitor cells at our endpoint is promising for putative long-term transplants. This proposes that these cells could have great potential for directed replacement therapies. While these advancements could be of great interest to the PD research community, we have to address several caveats. First, we only have established this protocol on benchmarked iPSC. But further in-depth characterization with more well-known mutations found in familial PD, such as GBA^E326K^, LRRK2^G2019S^ or the triplicate SNCA, could elucidate the validity of this enhanced protocol. Furthermore, the use of human embryonic stem cells has not been proposed for this study. More *in vitro* studies would be of interest to validate further this new protocol. Another caveat to address is the pre-clinical aspect, as the *in vivo* experiments were only used as proof-of-concept for therapeutical approaches. Further grafting experiments with PD-mutated hiPSC cell lines and a more in-depth preclinical analysis with Good Manufacturing Practice cell lines could detail if this protocol would be a much better advantage that the current available protocols, which so far have shown to be safe in clinical trials [12,13]. Moreover, the longevity of the grafting experiment was decided upon stable motor recovery, so we do not have an overview of the long-term effects of these grafts, whether other contaminants can appear or tumorigenesis in aged individuals.

## Conclusion

Summarizing, we have developed a multiomic-guided differentiation protocol which addresses the major challenge of subtype specificity in generating human mDA neurons relevant to PD. By leveraging single-cell genomics and epigenomics to identify and correct insufficient ventral patterning, we achieved robust enrichment of the PD-vulnerable SOX6⁺ A9 subtype, which yielded neurons with enhanced molecular fidelity, functional maturity, and the capacity to restore motor function upon transplantation.

Our work provides a robust platform for generating more accurate human cellular models of PD and establishes a foundational strategy for developing next-generation, subtype-enriched cell replacement therapies.

## Data Availability

Single-nuclei RNA-seq and ATAC-seq raw data have been deposited at EGA (EGA accession number: [EGAD50000002054](https://ega-archive.org/dacs/EGAC50000000805)). Key Resource Table (including Material RRID, protocol.io links for protocols used, accession numbers for sequencing datasets), microscopy data and images have been deposited at Zenodo at https://doi.org/10.5281/zenodo.17099441 and are publicly available as of the date of publication. All reused code used for this paper has been deposited at Github and linked to Zenodo at https://doi.org/10.5281/zenodo.17099441 and is publicly available as of the date of publication. This paper analyzes existing, publicly available data, accessible at https://doi.org/10.1038/s41593-022-01061-1; https://doi.org/10.1126/science.adf1226; https://doi.org/10.1126/science.add7046. Any additional information required to reanalyze the data reported in this paper is available from the lead contact upon request.

## Abbreviations

6-OHDA: 6-hydroxidopamine
BDNF: brain derived neurotrophic factor
DANs: dopaminergic neurons
EN1: engrailed-1
GDNF: glial derived neurotrophic factor
GO: gene ontology
hPSC: human pluripotent stem cell
iPSC: induced pluripotent stem cell
mDA: midbrain dopaminergic
mFP: midbrain floor plate
NbM: medial neuroblasts
PCR: polymerase chain reaction
PD: Parkinson’s Disease
ProgFP: floor plate progenitors
SCENIC: single-cell regulatory network inference and clustering
SHH: Sonic Hedgehog
SNpc: substantia nigra pars compacta
TF: transcription factor
UMAP: uniform manifold approximation and projection

## Acknowledgements

The authors acknowledge support from the National Genomics Infrastructure in Stockholm funded by Science for Life Laboratory, the Knut and Alice Wallenberg Foundation and the Swedish Research Council, and NAISS/Uppsala Multidisciplinary Center for Advanced Computational Science for assistance with massively parallel sequencing and access to the UPPMAX computational infrastructure. We would like to thank Eneritz Agirre for assisting with the deposit of snRNA-seq datasets. We are grateful for the assistance of the Molecular Neurobiology unit within the Department of Medical Biochemistry and Biophysics at the Karolinska Institute. We would like to posthumously thank Prof. Ernest Arenas, without whom this project would not have been possible.

## Ethics declarations

### Contributions

R.G.S., G.L. and E.A. conceptualized the experiments. R.G.S. and G.L. redacted the manuscript. R.G.S. performed *in vivo* experiments; R.G.S., G.L., R.K., C.S., K.S., C.T. and A.X. contributed to the *in vitro* protocol development and validation. J.C.K. contributed to the calcium imaging experiment. W.P. contributed to the HPLC analysis. R.G.S., G.L. and A.X. contributed to the *in vitro* transcriptional analysis; R.G.S. and C.A. performed the graft nuclei isolation, and R.G.S. and G.L. the *in vivo* transcriptional analysis. C.S., G.C.B., P.S. and O.D. contributed to supervision and reviewed the final manuscript.

---

**Extended Fig1.**
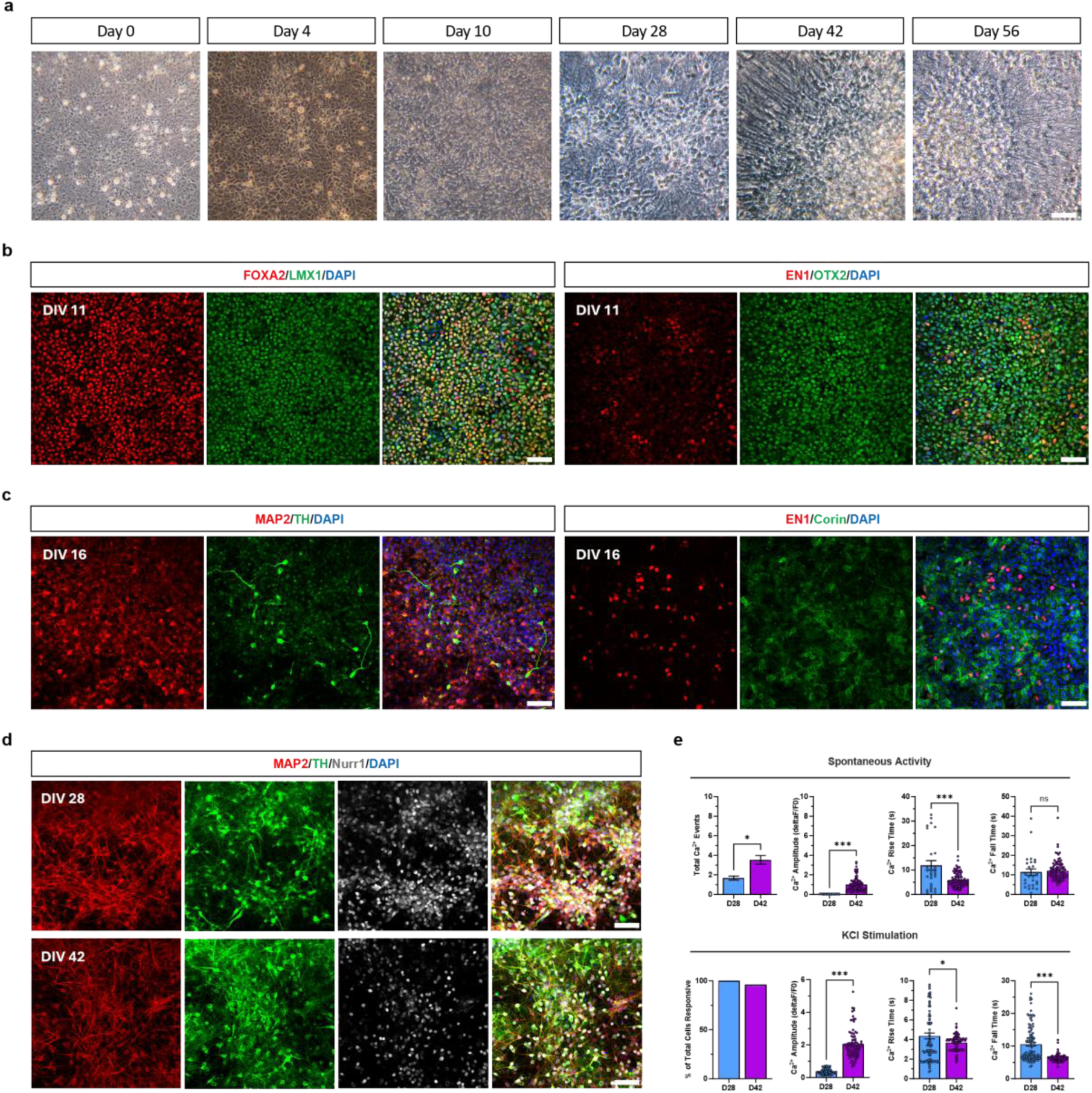
Chronological characterization of midbrain dopaminergic neurons derived from human induced pluripotent stem cell lines. a, Brightfield images showcase the sequential progression of neuronal morphology in mDA neurons. Representative immunocytochemical images showing **b,** staining of canonical transcription factors for midbrain floor plate patterning in the process of *in vitro* differentiation at DIV 11 and **c,** 16. **d**, Maturation of midbrain dopaminergic neurons at DIV 28 and 42. **e,** Calcium imaging recordings of spontaneous (top) and KCl-induced (bottom) activities on DIV 28 and 42, presenting the total number of calcium events per neuron recorded, and the percentages of cells responding to KCl-induced depolarization. Metrics of activities, including amplitude, rise time, and fall time of calcium signals, are depicted in the subsequent bar plots. White scale bars = 50 µm

**Extended Fig2.**
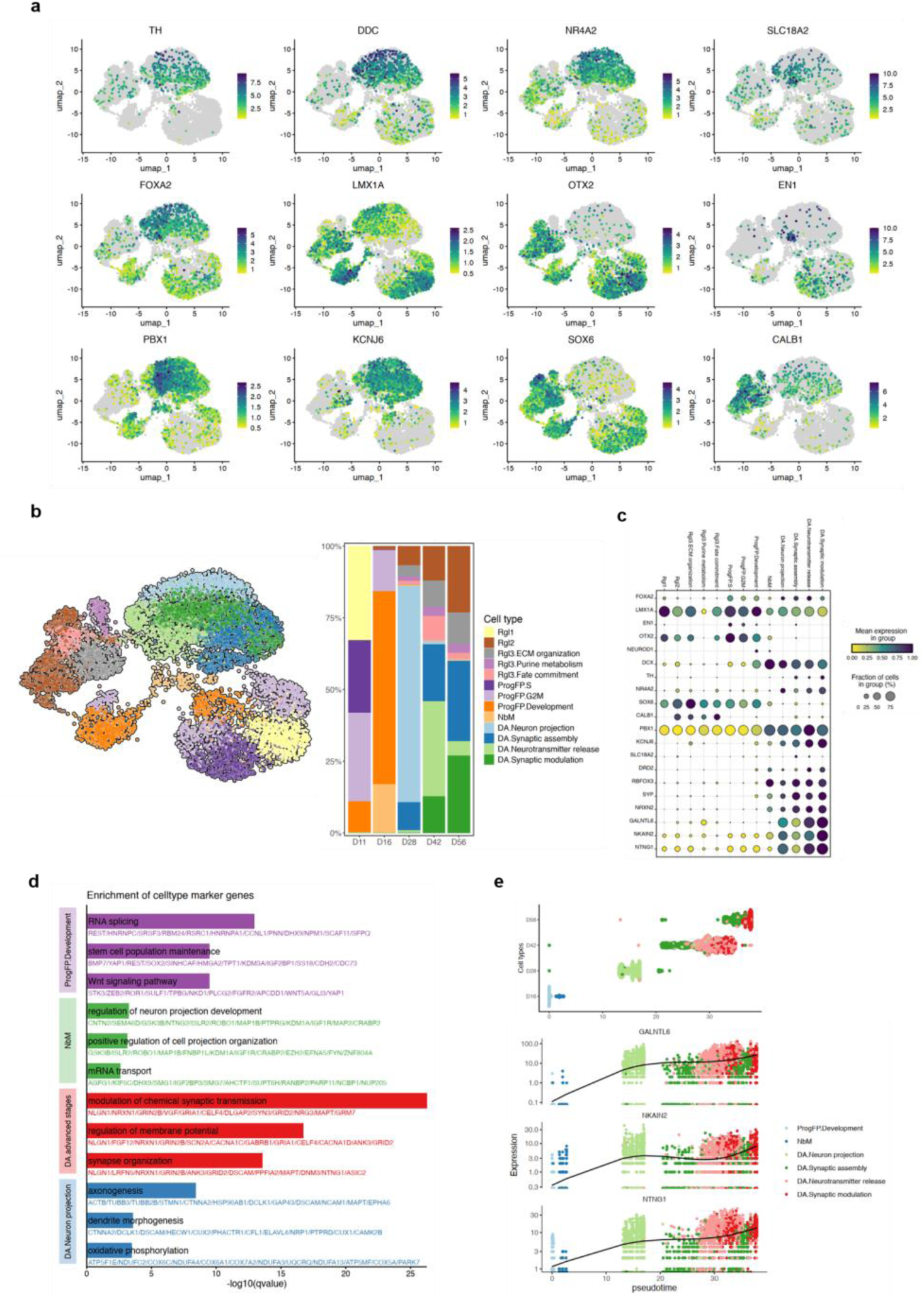
Multiomic single-cell analysis of human iPSCs at different stages of differentiation. a, Single-cell gene expression of classic dopaminergic marker genes (TH, DDC, NR4A2, and SLC18A2), midbrain regionalization marker genes (FOXA2, LMX1A, OTX2, and EN1), and subtyping marker genes (PBX1, KCNJ6, SOX6, and CALB1) across cells during dopaminergic neuron differentiation on the UMAP projection. **b**, Proportion of cells belonging to each cell type at each differentiation time point (left), and proportion of cells sub-setted by functionality (right). Detailed cell types were manually annotated with top-ranked marker genes and their enriched GO terms. **c**, Dot plot visualizing the expression levels and the percentage of cells expressing curated key genes across cell types during mDA neuron differentiation in detailed annotations. **d,** Gene Ontology (GO) enrichment analysis of mDA-lineage cells, showing distinct and related biological processes enriched in different cell types. **e,** Expression trends of selected genes along the pseudotime axis, these genes participate in specific biological processes pertinent to neuron maintenance or function.

**Extended Fig3.**
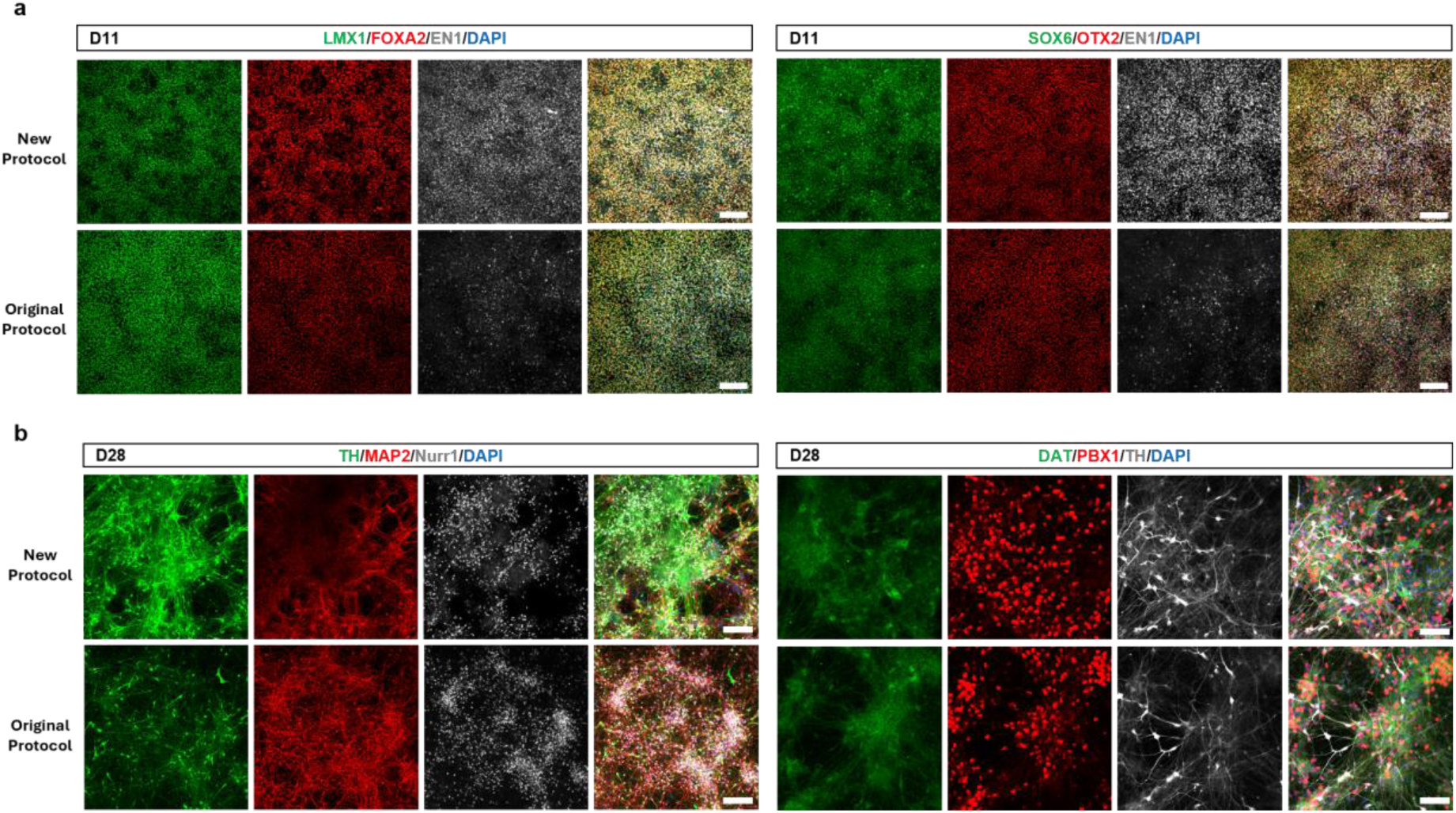
Further characterization of midbrain dopaminergic patterning of hiPSC comparing different protocols. a, Representative immunocytochemical images showing staining of canonical transcription factors for midbrain floor plate patterning of *in vitro* differentiation at DIV 11. Of note, SOX6 and EN1 presented higher intensity in the new protocol. **b**, Representative immunocytochemical images of markers for the maturation of midbrain dopaminergic neurons at DIV 28, where we can observe a higher amount of TH intensity in the new protocol. White scale bars = 100 µm

**Extended Fig4.**
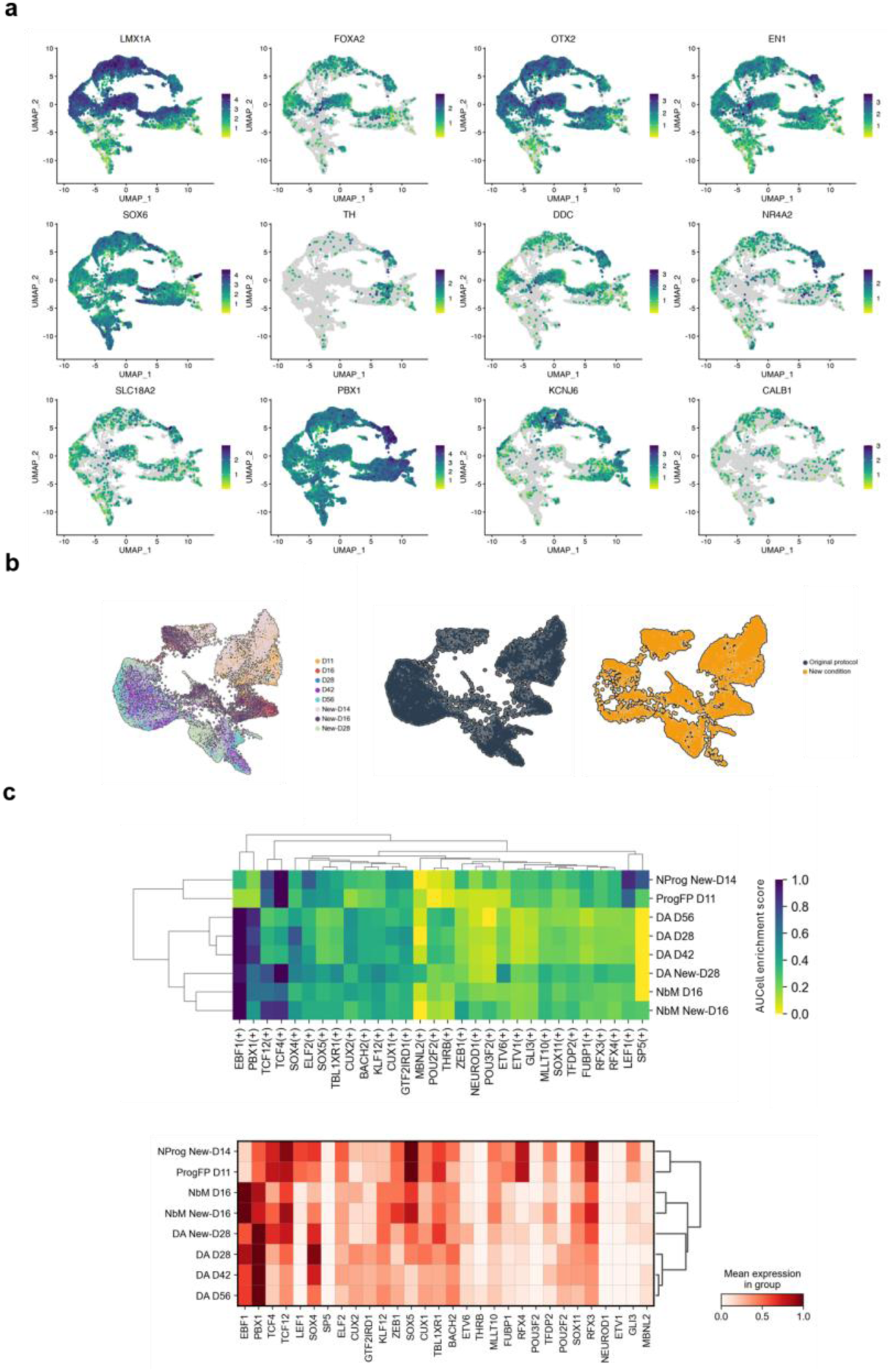
Single-nuclei analysis of midbrain cells generated from the new protocol.a, Single-nuclei gene expression of midbrain regionalization marker genes (LMX1A, FOXA2, OTX2, and EN1), classic dopaminergic marker genes (TH, DDC, NR4A2, and SLC18A2), and subtyping marker genes (PBX1, KCNJ6, SOX6, and CALB1) across cells during dopaminergic neuron differentiation on the UMAP projection. **b**, Integrated UMAP embedding of mDA differentiation in vitro using KOLF2.1J iPSCs, with nuclei from both the original (Figure 1) and new protocols (Figure 3). **c**, Heatmap of regulon activities (top) and expression levels of top 30 transcription factors (bottom) across selected mDA lineage cell types after the integration.

**Extended Fig5.**
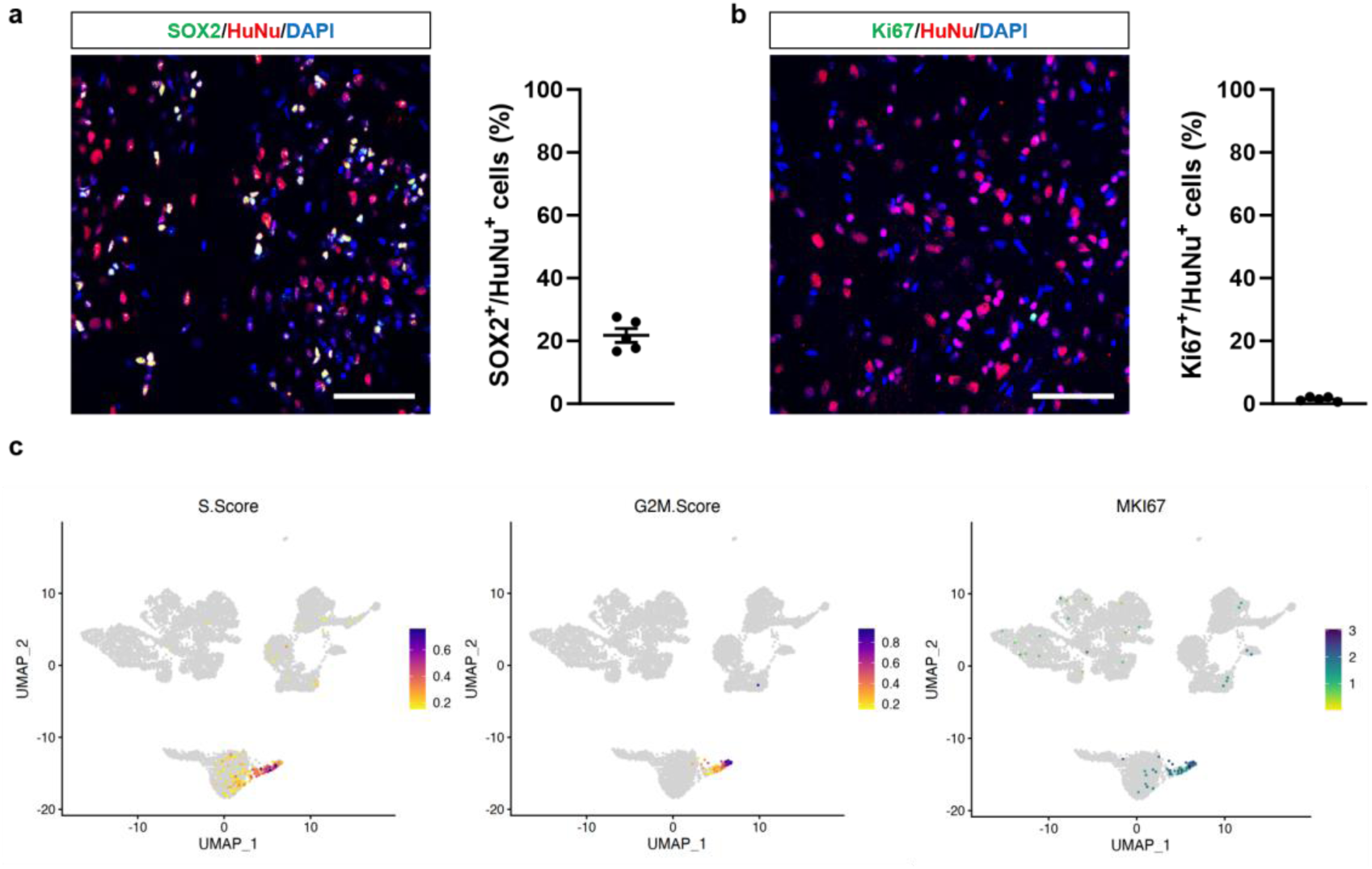
Lack of cell proliferation within grafted human DA progenitor cells. Immunohistochemical analysis of progenitor cells in human xenografts by **a**, SOX2 staining and **b**, active proliferation by Ki67 staining in all human nuclei (HuNu). Of note, no Ki67+ were found to be human. **c**, Single-nuclei analysis of active proliferative cells by S phase scoring, G2M phase scoring and expression of Ki67. Interestingly, proliferative cells were found in the oligodendrocyte lineage cluster. Bar scale in A,B = 50 µm. Each dot represents each graft, error bar = SEM.

**Extended Fig6.**
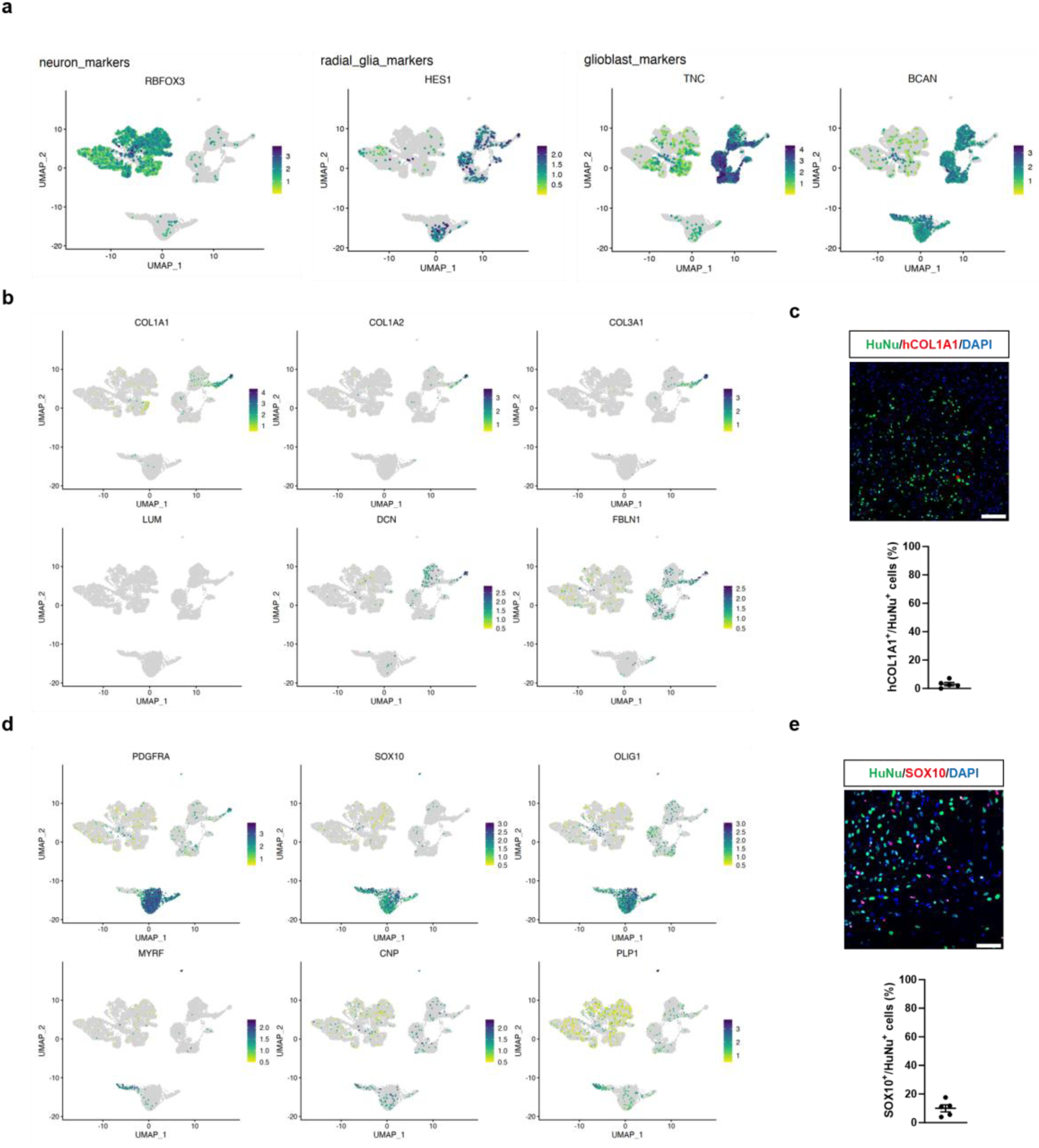
Further identity of cells by single-nuclei marker expression in *in vivo* xenografted cells. **a**, UMAP representation of top neuronal markers (RBFOX3), radial glia markers (HES1), and glioblasts (TNC, BCAN1). **b**, UMAP representation of expression of VLMC markers (COL1A1, COL1A2, COL3A1, LUM, DCN, FLBN1). **c**, Representative micrograph of immunohistochemistry of human grafted cells against human VLMC marker hCOL1A1 (top) with quantification of human cells positive for hCOL1A1 (bottom). **d**, UMAP representation of oligodendroglial markers (PDGFRA, SOX10, OLIG1, MYRF, CNP, PLP1). **e**, Representative micrograph of immunohistochemistry of human grafted cells against OPC marker SOX10 (top) with quantification of human cells positive for SOX10 (bottom). Scale bar = 100 µm.

**Extended Fig7.**
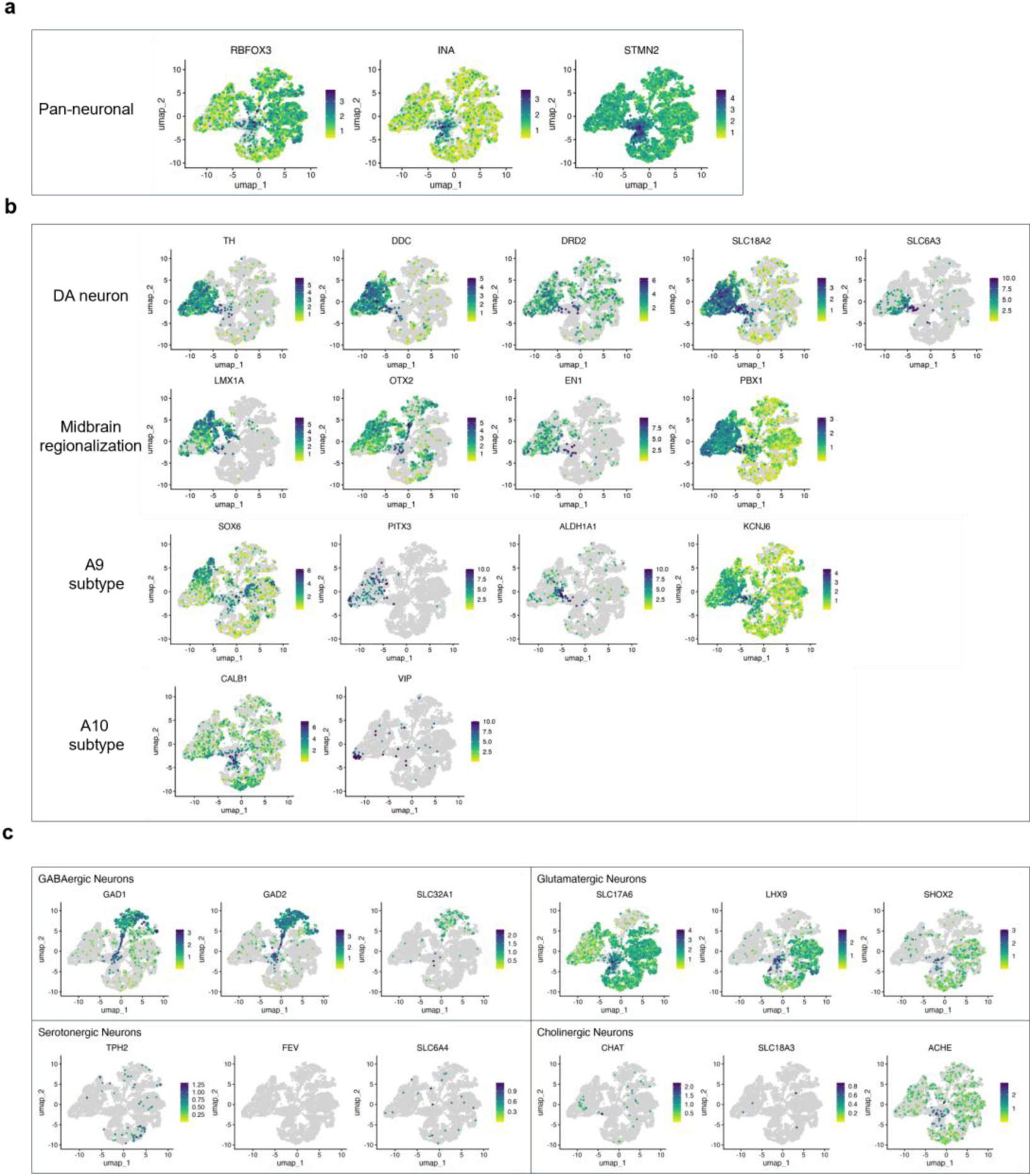
Identification of specific markers within the neuronal cluster of the human grafted cells. a, UMAP representation of pan-neuronal markers (RBFOX3, INA, STMN2). **b,** UMAP representation of DA neurons specificity based on DA neuronal general markers (TH, DDC, DRD2, SLC18A2, SLC6A3), for midbrain regionalized DA neurons (LMX1A, OTX2, EN1, PBX1), for A9 subtype DA neurons (SOX6, PITX3, ALDH1A1 and KCNJ6), and A10 subtype (CALB1, VIP). **c,** UMAP representation of non-DAergic neurons such as GABAergic neurons (GAD1, GAD2, SLC32A1), glutamatergic neurons (SCL17A6, LMX3, SHOX2), serotonergic neurons (TPH2, FEV, SLC6A4) and cholinergic neurons (CHAT, SLC18A3, ACHE). Of note, the grafted neuronal cluster lacks cells identifying as serotonergic neurons.

**Extended Fig8.**
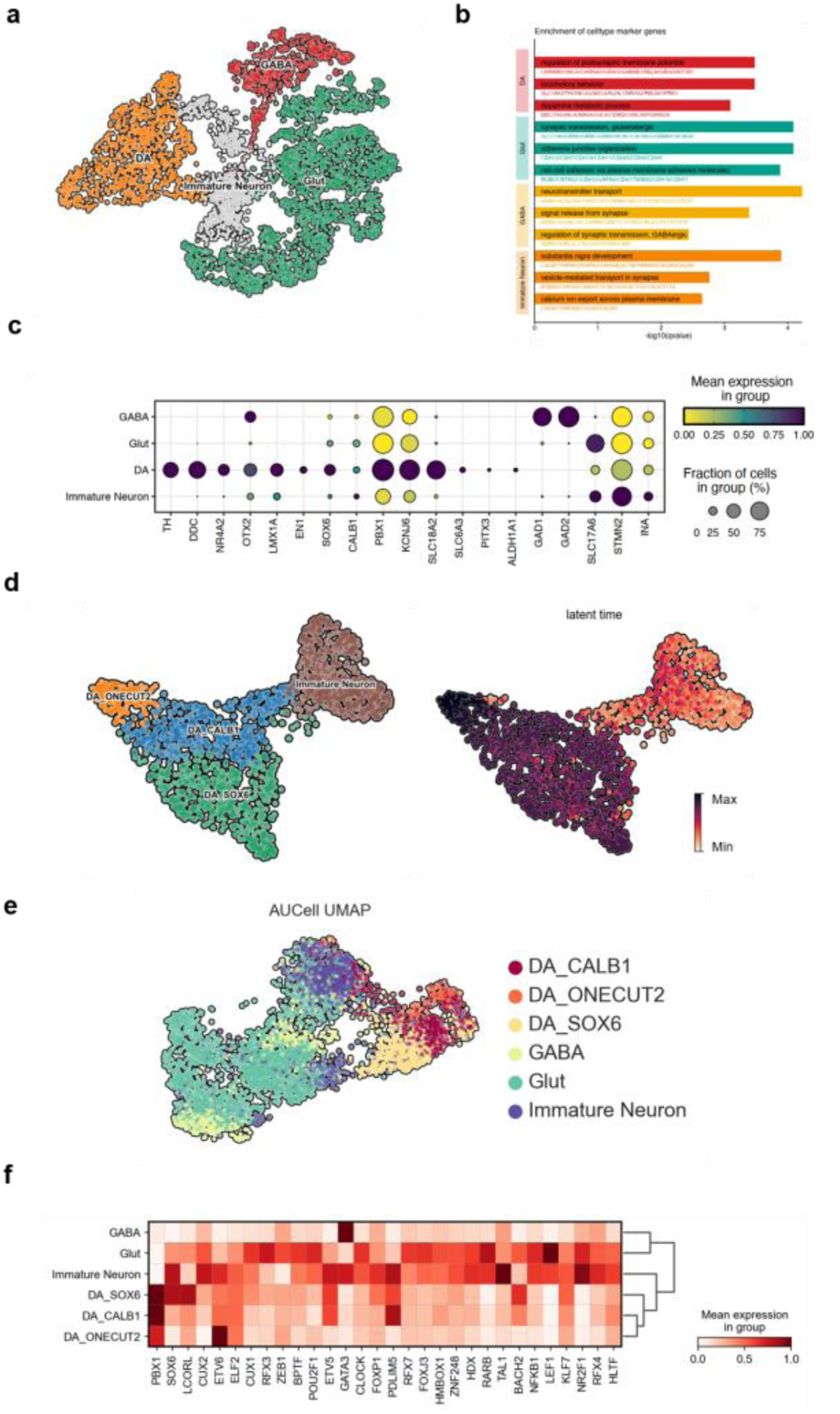
Single-nuclei analysis of neuronal clusters of *in vivo* xenograft. **a**, Annotated UMAP embeddings of neuronal population, including immature neurons, GABAergic neurons, glutamatergic neurons and dopaminergic neurons. **b**, Gene Ontology (GO) enrichment analysis of different types of neurons, showing distinct biological processes enriched according to their specific marker genes. **c,** Dot plot showing average gene expression and percentage of expressing cells for selected marker genes across annotated neuronal cell types. **d**, Visualization with UMAP embeddings showing mDA-related cell types, denotated in their subtypes and predicted latent time. **e**, UMAP embeddings of AUCell scores across different subtypes of neurons. **f**, Heatmap of expression levels of top 30 specific transcription factors across selected cell types.

