## Supplemental Figures for "Multi-omic guided iPSC dopaminergic differentiation leads to selective enrichment of PD-vulnerable SOX6+ midbrain neurons with motor recovery of PD models after transplantation"

### Slide 1
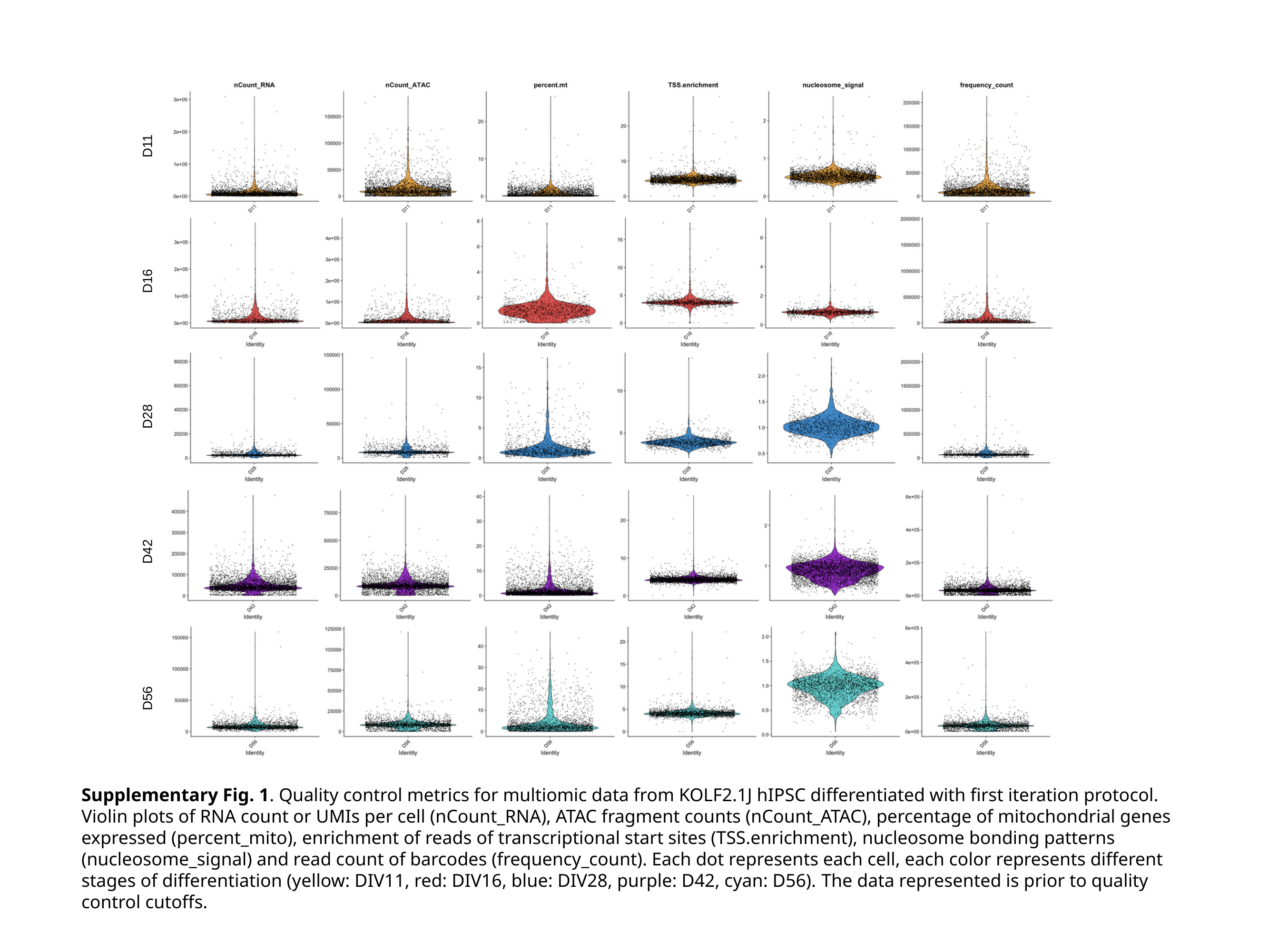

D11
D16
D28
D42
D56
Supplementary Fig. 1. Quality control metrics for multiomic data from KOLF2.1J hIPSC differentiated with first iteration protocol. Violin plots of RNA count or UMIs per cell (nCount_RNA), ATAC fragment counts (nCount_ATAC), percentage of mitochondrial genes expressed (percent_mito), enrichment of reads of transcriptional start sites (TSS.enrichment), nucleosome bonding patterns (nucleosome_signal) and read count of barcodes (frequency_count). Each dot represents each cell, each color represents different stages of differentiation (yellow: DIV11, red: DIV16, blue: DIV28, purple: D42, cyan: D56). The data represented is prior to quality control cutoffs.

### Slide 2
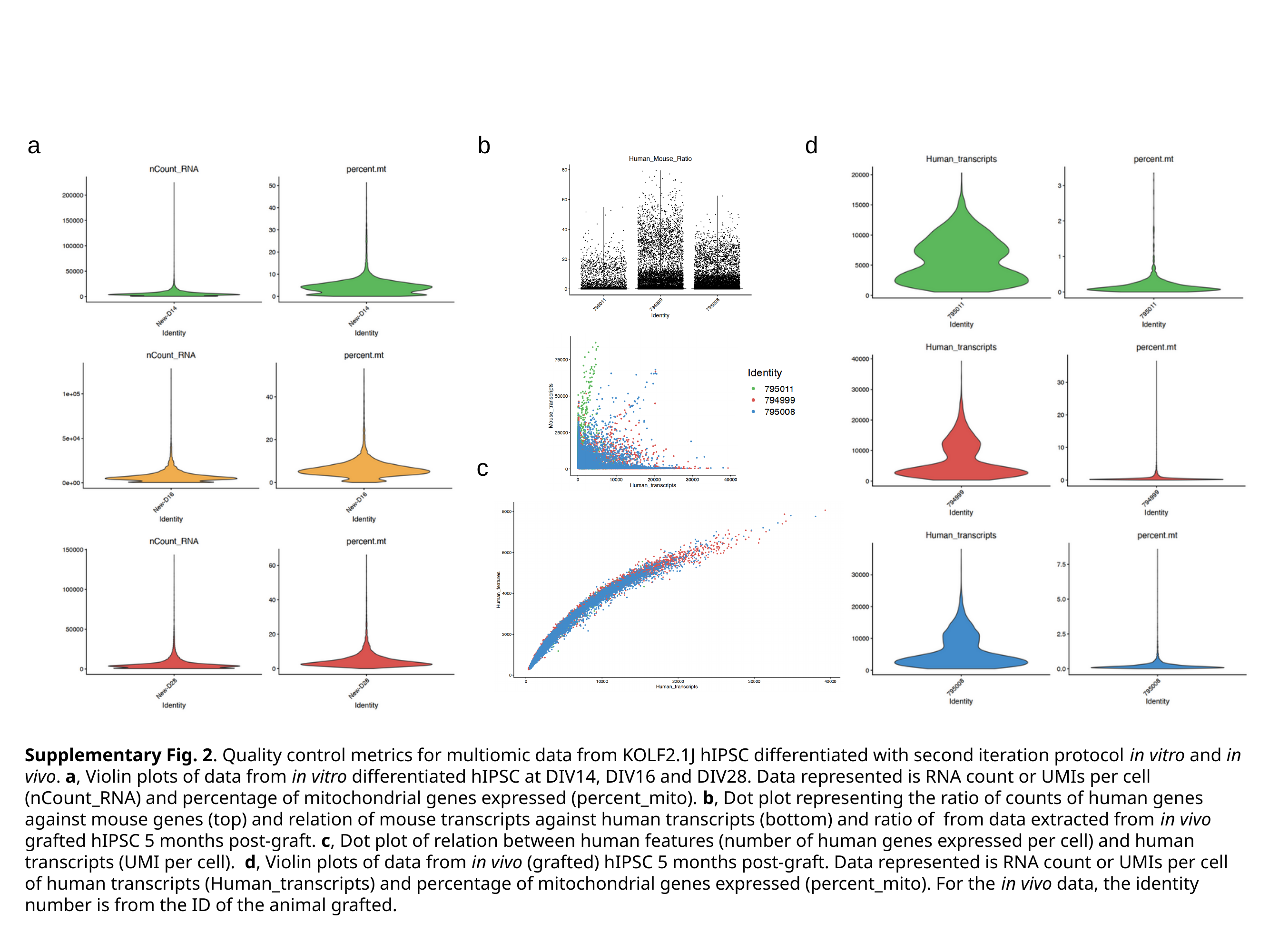

a
b
d
c
Supplementary Fig. 2. Quality control metrics for multiomic data from KOLF2.1J hIPSC differentiated with second iteration protocol in vitro and in vivo. a, Violin plots of data from in vitro differentiated hIPSC at DIV14, DIV16 and DIV28. Data represented is RNA count or UMIs per cell (nCount_RNA) and percentage of mitochondrial genes expressed (percent_mito). b, Dot plot representing the ratio of counts of human genes against mouse genes (top) and relation of mouse transcripts against human transcripts (bottom) and ratio of from data extracted from in vivo grafted hIPSC 5 months post-graft. c, Dot plot of relation between human features (number of human genes expressed per cell) and human transcripts (UMI per cell). d, Violin plots of data from in vivo (grafted) hIPSC 5 months post-graft. Data represented is RNA count or UMIs per cell of human transcripts (Human_transcripts) and percentage of mitochondrial genes expressed (percent_mito). For the in vivo data, the identity number is from the ID of the animal grafted.

### Slide 3
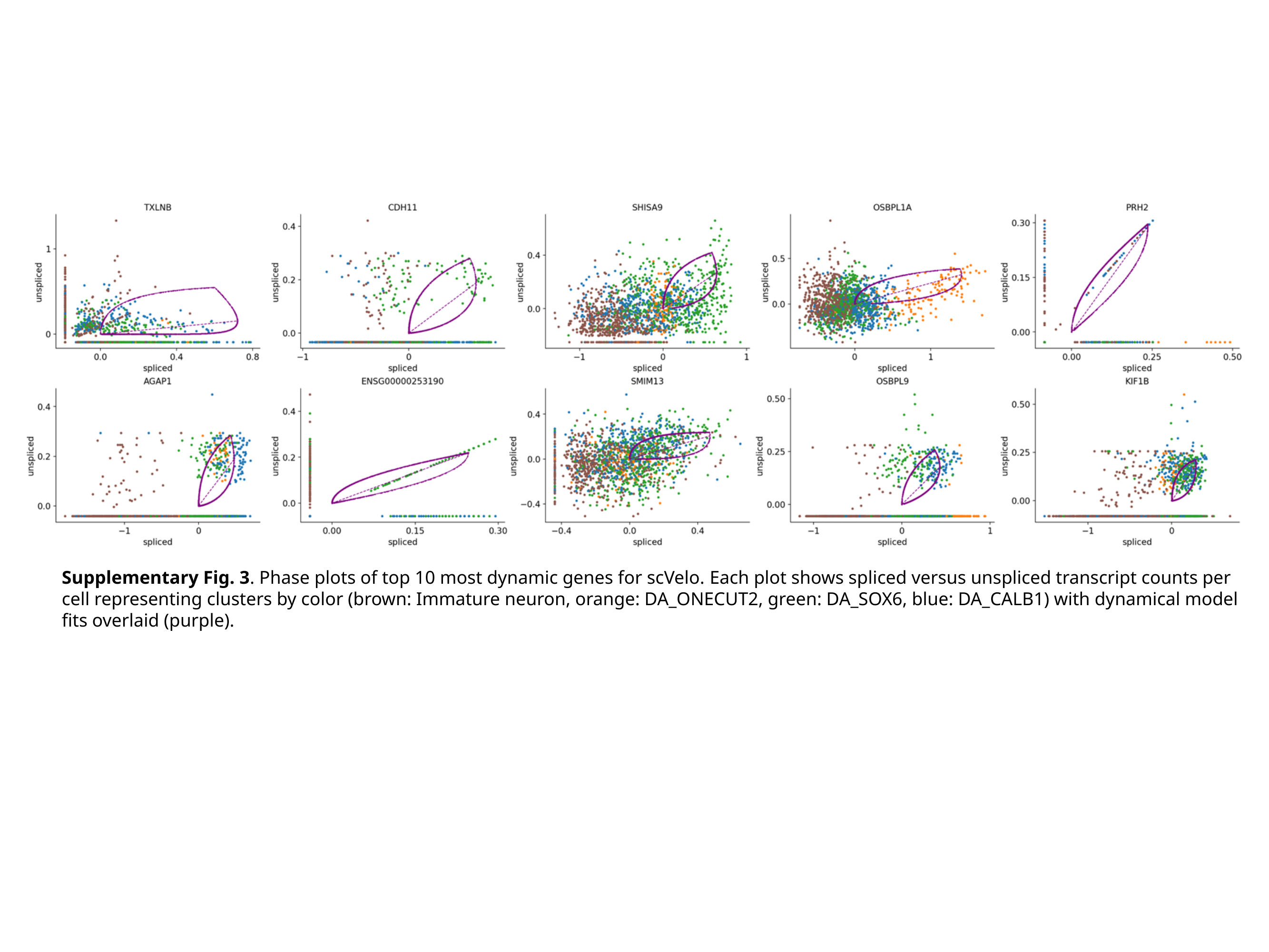

Supplementary Fig. 3. Phase plots of top 10 most dynamic genes for scVelo. Each plot shows spliced versus unspliced transcript counts per cell representing clusters by color (brown: Immature neuron, orange: DA_ONECUT2, green: DA_SOX6, blue: DA_CALB1) with dynamical model fits overlaid (purple).
